# Genome-wide, bidirectional CRISPR screens identify mucins as critical host factors modulating SARS-CoV-2 infection

**DOI:** 10.1101/2021.04.22.440848

**Authors:** Scott B. Biering, Sylvia A. Sarnik, Eleanor Wang, James R. Zengel, Varun Sathyan, Xammy Nguyenla, Erik Van Dis, Carmelle Catamura, Livia H. Yamashiro, Adam Begeman, Jessica C. Stark, D. Judy Shon, Douglas M. Fox, Andreas S. Puschnik, Carolyn R. Bertozzi, Jan E. Carette, Sarah A. Stanley, Eva Harris, Silvana Konermann, Patrick D. Hsu

## Abstract

SARS-CoV-2 can cause a range of symptoms in infected individuals, from mild respiratory illness to acute respiratory distress syndrome. A systematic understanding of the host factors mediating viral infection or restriction is critical to elucidate SARS-CoV-2 host-pathogen interactions and the progression of COVID-19. To this end, we conducted genome-wide CRISPR knockout and activation screens in human lung epithelial cells with endogenous expression of the SARS-CoV-2 entry factors ACE2 and TMPRSS2. These screens uncovered proviral and antiviral host factors across highly interconnected host pathways, including components implicated in clathrin transport, inflammatory signaling, cell cycle regulation, and transcriptional and epigenetic regulation. We further identified mucins, a family of high-molecular weight glycoproteins, as a prominent viral restriction network. We demonstrate that multiple membrane-anchored mucins are critical inhibitors of SARS-CoV-2 entry and are upregulated in response to viral infection. This functional landscape of SARS-CoV-2 host factors provides a physiologically relevant starting point for new host-directed therapeutics and suggests interactions between SARS-CoV-2 and airway mucins of COVID-19 patients as a host defense mechanism.

## INTRODUCTION

Severe acute respiratory syndrome coronavirus 2 (SARS-CoV-2) is a positive-sense RNA virus belonging to the *Betacoronavirus* genus within the *Coronaviridae* family (da Costa et al., 2020; Hartenian et al., 2020). The *Coronaviridae* family contains many human respiratory pathogens, from common cold coronaviruses to SARS-CoV-1 and Middle Eastern respiratory syndrome coronavirus (MERS-CoV) (da Costa et al., 2020; Cui et al., 2018; Hartenian et al., 2020). SARS-CoV-2 is the causative agent of coronavirus disease 2019 (COVID-19), and symptomatic cases of COVID-19 manifest with a diverse set of clinical outcomes ranging from fever and flu-like symptoms in non-severe cases to acute lung injury (ALI) and acute respiratory distress syndrome (ARDS) in severe cases (Blanco-Melo et al., 2020; Yang et al., 2020).

The SARS-CoV-2 viral life cycle begins with viral entry into susceptible angiotensin-converting enzyme II (ACE2)-expressing cells. After binding to the ACE2 receptor, the SARS-CoV-2 spike (S) glycoprotein is primed through a proteolytic cleavage event that can be mediated by the transmembrane serine protease 2 (TMPRSS2) on the cell surface, or in its absence, by the lysosomal endopeptidase cathepsin L (CTSL) following clathrin-mediated endocytosis. Subsequently, the primed S glycoprotein primarily mediates virus-cell fusion at the plasma membrane (TMPRSS2) or within an endosomal compartment (CTSL) (Trougakos et al., 2021; V’kovski et al., 2020; Zhu et al., 2021) The TMPRSS2-mediated cell surface entry route is believed to be dominant for SARS-CoV-2 in lung epithelial cells, as inhibition of TMPRSS2, but not CTSL, in primary lung epithelial cells is sufficient to inhibit viral infection (Hoffmann et al., 2020). Further, ACE2 and TMPRSS2 are largely co-expressed by the main targets of SARS-CoV-2 *in vivo*, such as epithelial cells within the lower and upper airway, the nasal passage, and the gut (Lukassen et al., 2020; Sungnak et al., 2020).

Following viral entry, the viral RNA genome is deposited into the host cytoplasm, where it undergoes translation. The resulting polyproteins are processed by viral and host proteases into an array of viral non-structural proteins (nsp). These proteins rearrange host membranes to form the endoplasmic reticulum-localized viral replication complex (RC). Safely sequestered within the RC, the SARS-CoV-2 genome is replicated by its RNA-dependent RNA polymerase (RdRP) complex. Newly produced viral genomes are then coated with the SARS-CoV-2 nucleoprotein (N) and egress through the secretory pathway, budding out of cells as new viral progeny ready to begin the cycle anew (Hartenian et al., 2020; V’kovski et al., 2020). During the SARS-CoV-2 lifecycle, the cell-intrinsic innate immune response can recognize viral pathogen-associated molecular patterns (PAMPs) and induce an antiviral cellular state. This typically involves activation of cell death programs, production of proinflammatory cytokines via the nuclear factor kappa-light-chain-enhancer of B cells (NF-κB) signaling pathway, and upregulation of type I interferons (IFN), which regulate expression of numerous interferon-stimulated genes (ISGs) (Katze et al., 2002; Yang et al., 2021). In turn, SARS-CoV-2 has evolved numerous strategies to antagonize or reroute these antiviral programs (Hartenian et al., 2020).

Each step of this viral life cycle requires a diverse array of host factors. A deeper understanding of the host-pathogen interactions of SARS-CoV-2 in permissive human lung cells will help illuminate molecular mechanisms of viral pathogenesis and host defense. Further, host-directed therapeutics hold great potential to alleviate COVID-19 disease burden, potentially posing less risk for viral escape compared to virus-targeted therapies. However, the host factors required for the SARS-CoV-2 viral life cycle in the human lung are largely unknown beyond a small number of critical entry factors.

While recent loss-of-function (LOF) screens have begun defining host factor requirements for SARS-CoV-2 infection, these studies employed host gene knockout either in non-epithelial cell lines or cell lines that do not endogenously express ACE2 and TMPRSS2 (Baggen et al., 2021; Bailey and Diamond, 2021; Daniloski et al., 2021; Schneider et al., 2021; Wang et al., 2021; Wei et al., 2021; Zhu et al., 2021). LOF screens remove host factors that mediate viral infection and cytopathic effect, allowing for the identification of proviral genes. In contrast, gain-of-function (GOF) screens can identify antiviral factors that enhance viral restriction upon upregulation. Performing both screens in a bidirectional manner can therefore illuminate host pathways with bimodal roles. Finally, as expression of TMPRSS2 defines the cell-entry route of SARS-CoV-2, distinct host dependency factors would be expected to emerge from cell lines with varying levels of this protease (Koch et al., 2020; Zhu et al., 2021).

We therefore set out to define the host-pathogen interactions required for facilitating or restricting SARS-CoV-2 infection in Calu-3 cells, which are human lung cells endogenously expressing both ACE2 and TMPRSS2. We conducted genome-scale loss-of-function and gain-of-function CRISPR screens to generate a systematic functional map of host dependency and host restriction factors. Pathway analysis and secondary validation of the screen hits revealed diverse cellular components involved in modulating cellular proliferation, intercellular junctional complexes, the cytoskeleton, inflammatory signaling, and mucin glycoproteins.

The gene hits identified in our bidirectional dataset are enriched in single cell RNA sequencing datasets of lung epithelial cells from healthy individuals and COVID-19 patients, underscoring their physiological relevance. We further validated a network of membrane-tethered mucins as antiviral factors, which are O-glycosylated proteins known to modulate bacterial infection and restrict influenza A virus infection (Lu et al., 2006; McAuley et al., 2017; Zanin et al., 2016). We demonstrate that these mucins restrict SARS-CoV-2 infection at the entry stage, suggesting that mucin expression in human lungs may modulate SARS-CoV-2 infection. Taken together, our bidirectional CRISPR screens dissect the proviral and antiviral roles of the densely interconnected genetic landscape of SARS-CoV-2 host factors, highlighting mucins as potential restriction factors of SARS-CoV-2 infection in COVID-19 patients.

## RESULTS

### Genome-wide bidirectional CRISPR screens reveal a functional map of SARS-CoV-2 host factors

Our LOF and GOF screens used virus-mediated cell death as a functional readout, exploiting Cas9 nucleases for gene ablation and dCas9 with functionalized guide RNA aptamers to recruit synergistic transcriptional activators for target upregulation **(Figure 1)** (Hsu et al., 2014). We utilized a human lung epithelial cell line (Calu-3) that is highly permissive to SARS-CoV-2 infection and exhibits dramatic cytopathic effect (CPE) without the need for exogenous overexpression of SARS-CoV-2 entry factors such as ACE2 and TMPRSS2. We first transduced Calu-3 cells with constructs encoding Cas9 nuclease (LOF) or the transcriptional activators dCas9-VP64 + PP7-P65-HSF1 (GOF), followed by lentiviral delivery of CRISPR guide libraries at low multiplicity of infection **(Figure 1A)** (Konermann et al., 2015; Sanson et al., 2018).

**Figure 1:**
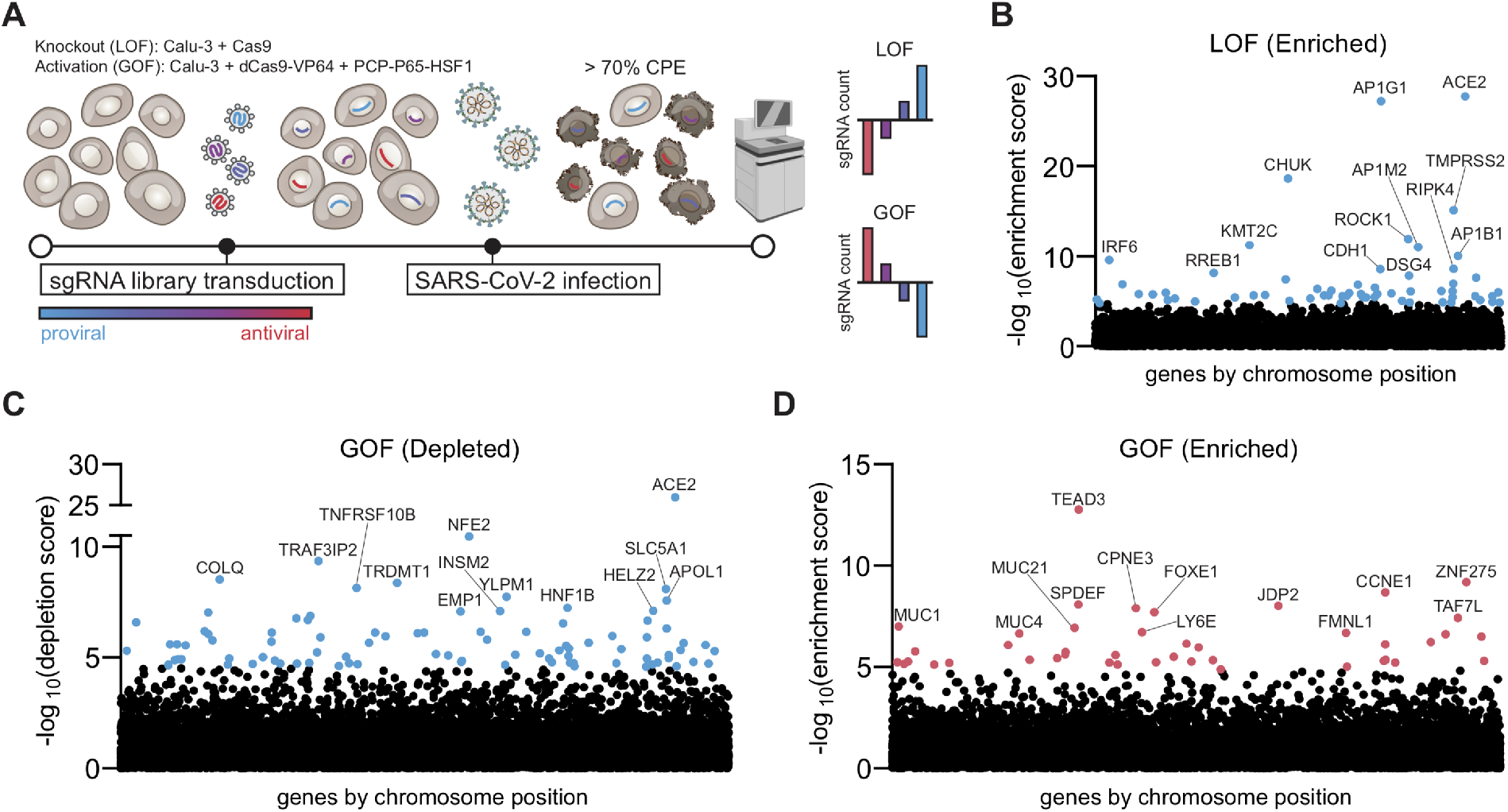
Bidirectional CRISPR screens identify host factors critical for SARS-CoV-2-mediated cytopathic effect. **A.** Schematic of genome-wide CRISPR knockout and activation screens for SARS-CoV-2 host factors, conducted in parallel. Calu-3 cells stably expressing Cas9 for the loss-of-function (LOF) screen or dCas9 and transcriptional activators for the gain-of-function (GOF) screen were transduced with pooled guide RNA libraries. Following infection with SARS-CoV-2, cells were harvested after at least 70% cytopathic effect was evident. Nextgeneration sequencing was performed to identify host factors and assign proviral and antiviral roles based on guide RNA enrichment or depletion compared to uninfected controls. **B.** Manhattan plot displaying the top 13 enriched genes identified in the LOF screen. **C.** Manhattan plot displaying the top 13 depleted genes in the GOF screen. **D.** Manhattan plot displaying the top 13 enriched genes in the GOF screen. All genes are ranked based on MAGeCK robust rank aggregation score. Red dots indicate putative antiviral genes with FDR<0.05, blue dots indicate putative proviral genes with FDR <0.05.

Both screens were conducted with four biological replicates maintaining greater than 1000X single guide RNA (sgRNA) coverage. Following SARS-CoV-2 infection with the USA/WA-1 isolate, genomic DNA was harvested after at least 70% of cytopathic effect (CPE) was observed. Nextgeneration sequencing of sgRNAs enabled identification of genes enriched and depleted relative to uninfected control cells **(Tables S1-S3)**. We interpreted guides targeting genes emerging from the LOF-enriched **(Figure 1B)** and GOF-depleted (**Figure 1C**) categories to be proviral, and guides targeting genes from the GOF-enriched category (**Figure 1D**) to be antiviral.

The top gene from both the LOF-enriched and GOF-depleted screens was ACE2, the SARS-CoV-2 cell-entry receptor, validating the phenotypic selection of the host factor screens. TMPRSS2 was also highly enriched in our LOF screen, suggesting that SARS-CoV-2 infection of Calu-3 cells is dependent on the dominant SARS-CoV-2 S glycoprotein-priming mechanism in lung epithelial cells (Hoffmann et al., 2020; Ou et al., 2021).

Beyond ACE2, our top proviral gene hits include AP1G1 and CHUK in the LOF-enriched analysis as well as NFE2 and TRAF3IP2 in the GOF-depleted direction **(Figure 1B, 1D)**. AP1G1 encodes a clathrin-adaptor protein potentially involved in clathrin-mediated intracellular trafficking, egress, or entry of SARS-CoV-2. CHUK encodes for IKK-alpha which is a critical component in regulation of the NF-κB signaling pathway, a transcriptional complex commonly activated by viral infections that regulates diverse events including proinflammatory cytokine production and cell survival (Santoro et al., 2003; Taniguchi and Karin, 2018). NFE2 is a transcription factor regulating diverse cellular responses, including cell differentiation, and has been shown to be upregulated in response to SARS-CoV-2 infection (Gasiorek and Blank, 2015; Thair et al., 2021). TRAF3IP2 encodes for ACT1, an adaptor protein possessing E3-ubiquitin ligase activity involved in IL-17 and NF-κB signaling pathways (Li et al., 2000). The role of NF-κB and IL-17 signaling may be directly proviral by establishing a cellular environment conducive to viral replication, or may negatively affect the host cell by promoting cell death (DeDiego et al., 2014; Kircheis et al., 2020; Park and Hong, 2016; Poppe et al., 2017; Su et al., 2020).

On the antiviral side, our GOF screen identified TEAD3, ZNF275, and CCNE1 as the top 3 enriched genes **(Figure 1D and Table S3)**. TEAD3 is a transcription factor involved in the TGF-β and Hippo signaling pathways, which can regulate cell proliferation (Ma et al., 2019). Hippo signaling in particular is activated by diverse stimuli including viral infection and is regulated through kinases such as the LOF-enriched hits TAOK1 and TAOK2 (Plouffe et al., 2016). CCNE1 encodes for cyclin E1, a regulatory subunit of cyclin-dependent kinase 2 (CDK2) which is required for the G1/S cell cycle transition (Honda et al., 2005). As numerous studies have reported cell cycle arrest as a requirement for optimal viral replication, enhanced cell proliferation resulting from CCNE1 overexpression is likely refractory to SARS-CoV-2 replication (Bagga and Bouchard, 2014; Davy and Doorbar, 2007; Fan et al., 2018). In fact, the SARS-CoV N protein has been previously shown to inhibit CCNE1, suggested to be a strategy of viral-mediated cell-cycle arrest to route cellular resources towards viral replication (Surjit et al., 2006). Finally, ZNF275 is a zinc finger protein that may be involved in transcriptional regulation (Haston et al., 2006). Taken together, viral entry and trafficking factors, components involved in proinflammatory responses, and cell proliferation regulators are critical determinants of SARS-CoV-2-mediated cell death in Calu-3 cells.

### Insight into SARS-CoV-2 host dependency factors, pathways, and interaction networks

To gain more systematic insight into critical proviral host pathways and interactions for SARS-CoV-2 infection, we conducted two analyses of our top 100 enriched and depleted hits from our LOF and GOF screens, respectively. First, we performed a protein-protein interaction network enrichment analysis to define putative interactions between our top 100 genes, clustering hits into distinct interaction clusters based on both direct and indirect connections (**Figure 2A** and **Figure S1A**). Next, we performed a gene set enrichment analysis to identify enriched biological pathways (**Figure 2B and Figure S1B**). The identified apoptotic signaling pathways are expected to be directly involved in mediating virally-induced cytopathic effects, while canonical interferon signaling may be enriched in the LOF screen through modulation of cell proliferation or cell death pathways. We also found components of NF-κB-mediated inflammatory signaling, cell-cell junctional complexes, cytoskeletal remodeling, adaptor-mediated clathrin transport, and cell cycle regulation as enriched in our proviral screens. The interaction network analysis further revealed specific interconnections between genes involved in these host pathways that mediate SARS-CoV-2 infection in Calu-3 cells **(Figure 2A and Figure S1B)**.

**Figure 2:**
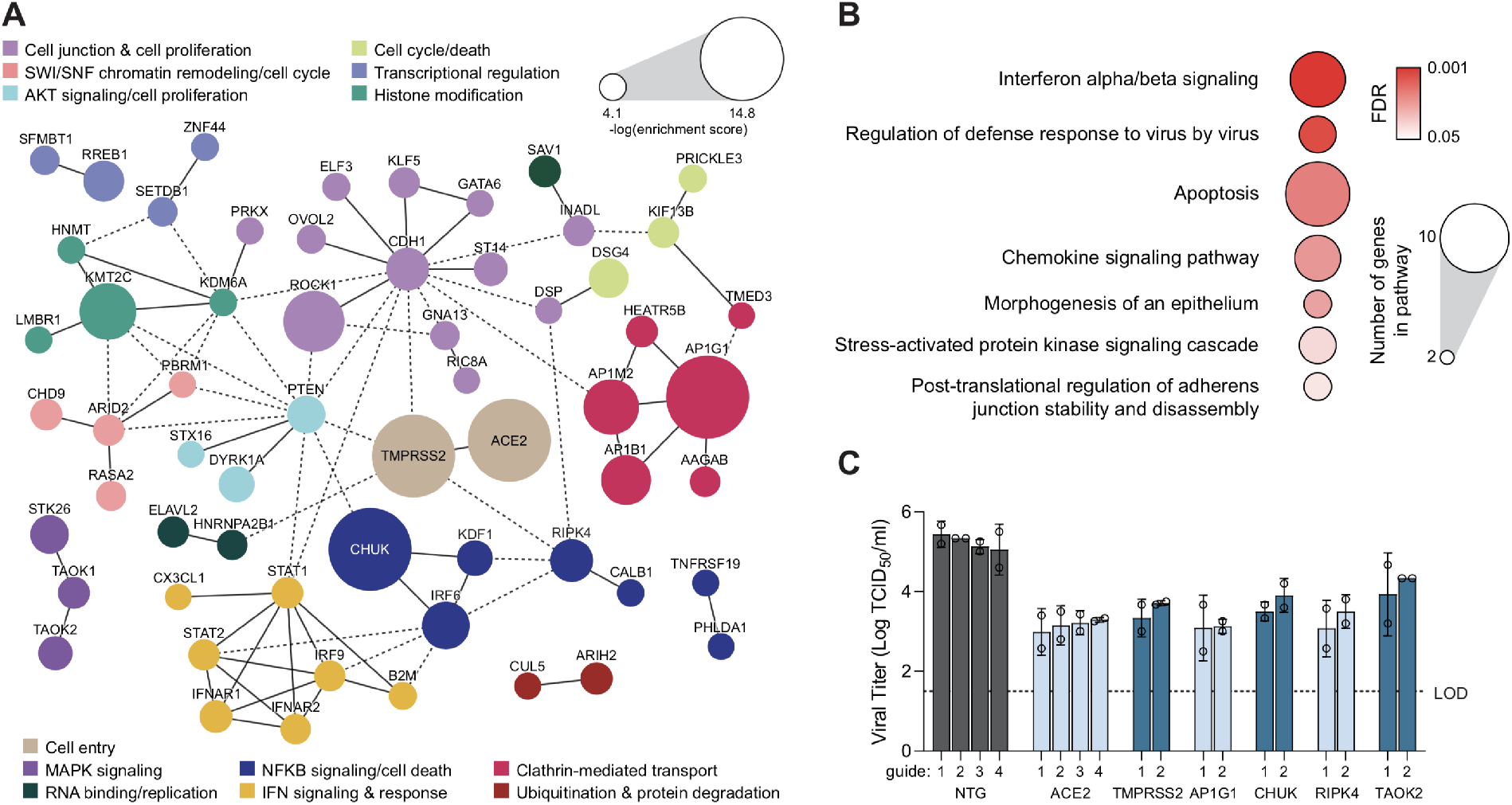
Host dependency factors and pathways of SARS-CoV-2 in lung cells revealed by genome-wide, loss-of-function screening. **A.** Protein-protein interaction network for top 100 enriched hits identified in the CRISPR LOF screen based on STRING analysis. Solid lines between genes indicate direct interaction, dashed lines indicate indirect connections. Nodes are color-coded by functional groups and scaled according to screen enrichment RRA score. **B.** Pathway analysis of top 100 enriched hits indicates significantly overrepresented pathways with putative proviral roles. Circle size indicates the number of genes within each pathway, color indicates FDR of pathway enrichment. **C**. Individual TCID_50_ validation of the effect of KO of six putative proviral hits on SARS-CoV-2 viral titer in Calu-3 infected at an MOI of 0.05 for 48 hours (input viral inoculum not washed away). Each gene was targeted with at least two separate guides. Error bars denote mean ± SEM, n = 2. Dotted line indicates the limit of detection (LOD) of the assay. NTG = non-targeting guide control.

To confirm the requirements of individual genes for SARS-CoV-2 infection, we generated individual Calu-3 CRISPR lines with two distinct guides per gene for 8 of our top LOF-enriched hits and GOF-depleted gene hits. These gene hits represent distinct pathways and interaction clusters predicted to be critical for SARS-CoV-2 infection including clathrin-adaptor proteins and NF-kB inflammatory responses (**Figure 2A)**. We observed reduced infectious virus production using a median tissue culture infectious dose (TCID_50_) assay from SARS-CoV-2 infected cells transduced with guides targeting ACE2 and TMPRSS2, as well as AP1G1, CHUK, RIPK4, and TAOK2, compared to control Calu-3s (**Figure 2C**). This supports a key role of the adaptor protein 1 (AP1) clathrin adaptor complex in the viral life cycle, of which we identified three subunits (AP1G1, AP1M2, AP1B1) as well as two direct interaction partners in our KO screen (HEATR5B, AAGAB) **(Figure 2A)**. The observed proviral effects of RIPK4 and CHUK suggest that certain components of NF-kB signaling are beneficial for SARS-CoV-2 infection and may be actively regulated by the virus to promote a viral replicative niche. It has been previously observed that NF-kB pathways can play both proviral and antiviral roles which can be actively regulated and rerouted by other coronaviruses and influenza A (Poppe et al., 2017; Schmolke et al., 2009).

For GOF-depleted screen validation, lentiviral transduction with CRISPR activation components confirmed that upregulation of ACE2, MEX3B, APOL1, and CDKN2B increased viral infection and replication compared to non-targeting (NTG) control cells, as expected by their depletion scores in the GOF screen (**Figure S1C**). CDKN2B encodes for the cyclin-dependent kinase inhibitor 2B, which controls the progression from G1 to S phase. Negative regulation of cell proliferation was the top pathway found among the depleted genes of the GOF screen (**Figure S1B**), supported by previous observations that the nucleocapsid protein (N) of SARS-CoV actively inhibits the functions of cyclin-dependent kinases (CDKs) to arrest the cell cycle (Surjit et al., 2006).

### Comparative analysis of LOF screens reveals unique SARS-CoV-2 dependencies in lung epithelial cells with endogenous entry factors

We next conducted a comparative analysis of our LOF screen hits with recent LOF studies of SARS-CoV-2 proviral host factors in different cell lines (Baggen et al., 2021; Daniloski et al., 2021; Schneider et al., 2021; Wang et al., 2021; Wei et al., 2021). Other than ACE2, we noticed surprisingly low overlap in the top-ranked hits (**Figure 3A**). Pairwise comparisons between our Calu-3 screen and other screens resulted in only 0-4 overlapping genes in the top 100 hits, suggesting that many of our hits are specific to the cell line we used. Calu-3 cells produce and respond to interferon and endogenously express ACE2 and TMPRSS2, in contrast to monkey kidney epithelial cells (Vero-E6) and human hepatocytes (Huh7.5) with defective type I IFN responses or entry factor-engineered cell lines (Huh7.5.1 overexpressing ACE2 and TMPRSS2, and A549 overexpressing ACE2) (Desmyter et al., 1968; Mosca and Pitha, 1986; Sumpter et al., 2005). We therefore hypothesized that our unique screen hits may be due to distinct properties of Calu-3 lung epithelial cells, such as TMPRSS2-dependent viral entry or functional inflammatory responses.

**Figure 3:**
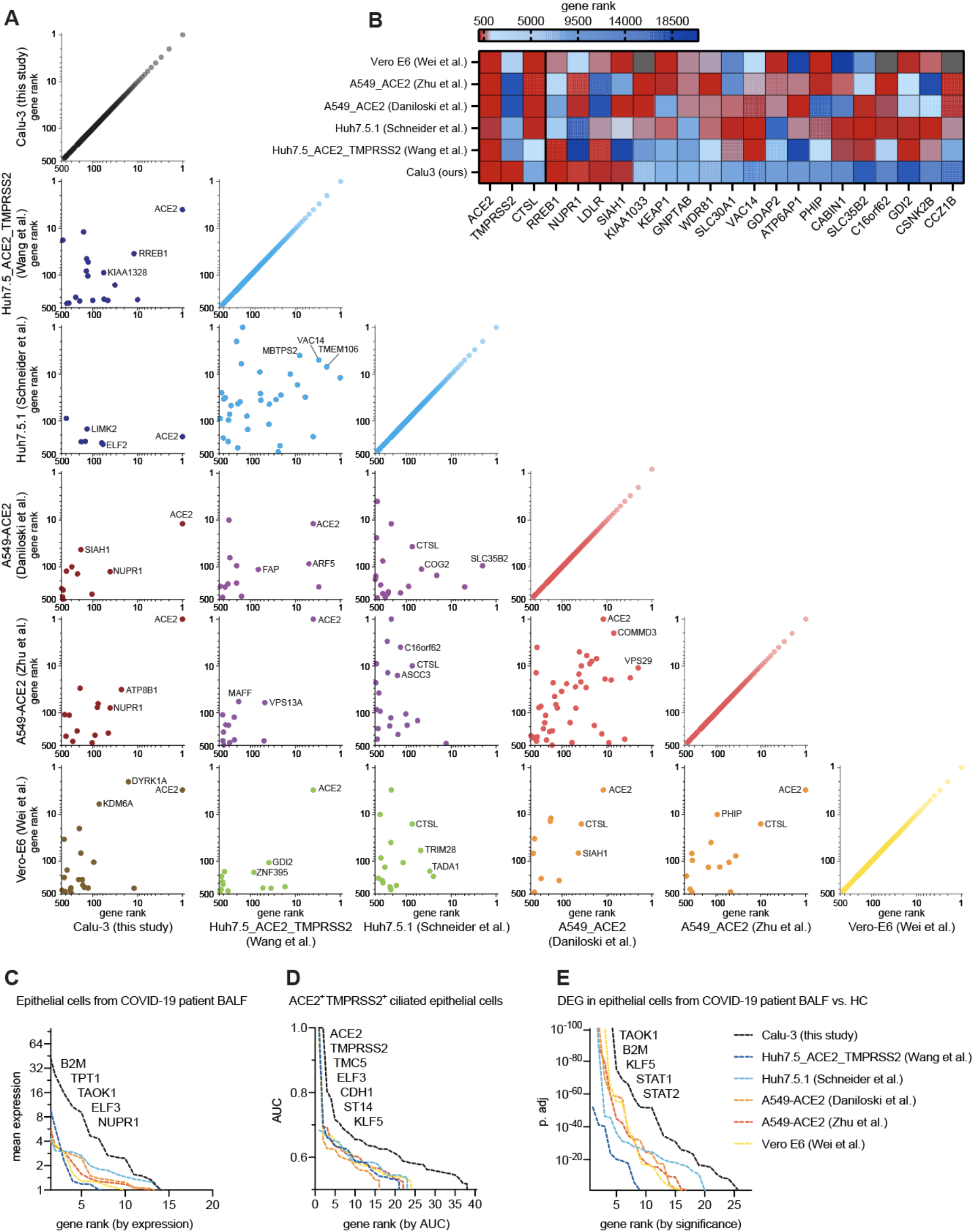
Comparative analysis of loss-of-function screens reveals cell type-specific host factor landscapes of SARS-CoV-2. **A.** Correlation plots of top 500 hits ranked by RRA score from this study and previously reported SARS-CoV-2 knockout screens performed in different cell lines. Plots are color-coded by cell line. Black: Calu-3 (human epithelial lung adenocarcinoma cell line), blue: Huh7.5 and Huh7.5.1 (human hepatocyte-derived cell lines), red: A549-ACE2 (ACE2-overexpressing human lung adenocarcinoma cell line), yellow: Vero-E6 (African Green Monkey kidney epithelial cell line). The top three overlapping genes with the lowest sum of rank position are displayed. **B**. Heat map indicating screen rank of key SARS-CoV-2 entry factors (left side) and all other top 500 ranked hits present in at least three screens (right side). **C.** Top 100 hits across LOF screens ranked by their expression levels in lung epithelial cells from COVID-19 patient BALF single cell RNA-seq data from Liao et al., 2020. **D.** Top 100 hits across LOF screens ranked by AUC value for ACE2^+^TMPRSS2^+^ ciliated human lung epithelial cells based on scRNA-seq meta-analysis, data from Muus et al., 2021. **E.** Top 100 hits across LOF screens ranked based on differential expression in lung epithelial cells from COVID-19 patients compared to healthy individuals, data from Liao et al., 2020.

Comparing across all screens, the cell type chosen for infection is the dominant factor determining screen hit overlap, rather than differences in viral strain, multiplicity of infection (MOI), or timeline of genomic DNA harvest. CRISPR knockout screens conducted by independent groups in the same cell lines, A549-ACE2 and Huh7 cells, had 21 and 13 overlapping genes in the top 100 hits, respectively. Strikingly, all other pairwise comparisons yielded 0-4 overlapping genes in the top 100 hits, as we had observed for our Calu-3 screen **(Figure 3A)**. The gene hits common to our screen and at least two other studies are involved in biological processes such as regulation of transcriptional responses (RREB1, NUPR1), cholesterol uptake (LDLR), and protein ubiquitination (SIAH1) **(Figure 3B)**. In contrast, a large cluster of gene hits enriched across at least 3 other screens - but not our Calu-3 screen - include genes encoding vacuolar-associated proteins important for endolysosomal trafficking (CTSL, ATP6AP1, GNPTAB, VAC14, WDR81, GDI2, and CCZ1B) **(Figure 3B)**. In contrast, members of the AP1 clathrin adaptor complex were uniquely identified in our screen (**Figure 2A**). These hits likely highlight differences in the CTSL- and TMPRSS2-dependent entry and trafficking routes of SARS-CoV-2. Taken together, host factors regulating endosomal maturation and CTSL function may be dispensable for SARS-CoV-2 infection when TMPRSS2 is present.

To assess the physiological relevance of our screen hits, we conducted comparative analyses of two distinct single cell RNA-seq studies: a dataset of human bronchoalveolar lavage fluid (BALF) from COVID-19 patients and controls (Liao et al., 2020), and a recent meta-analysis of healthy human lung cell types (Muus et al., 2021). First, we identified the gene expression profiles from total lung epithelial cells of COVID-19 patients. Of the top 100 ranked genes within each screen, a greater fraction of our Calu-3 screen hits was expressed at higher levels compared to gene hits from other screens (**Figure 3C**), including genes involved in inflammatory responses (B2M, TAOK1) and cell proliferation (ELF3, TPT1, and NUPR1).

Next, we analyzed ciliated epithelial cells co-expressing ACE2 and TMPRSS2, presumed to be primary cellular targets of SARS-CoV-2 infection in humans (He et al., 2020; Ravindra et al., 2021; Sajuthi et al., 2020; Sungnak et al., 2020). We again observed greater enrichment of our Calu-3 gene hits compared to gene hits from other screens, implying Calu-3 cells may be more representative of physiologically relevant ACE2^+^/TMPRSS2^+^ cells in human lungs compared to cell lines employed in other screens (**Figure 3D**). Top gene hits found in our Calu-3 screen that were specific to ACE2^+^/TMPRSS2^+^ ciliated lung cells include regulators of cell proliferation and migration (TMC5, ST14, KLF5) or cell-cell adhesion (CDH1). Finally, comparing gene expression profiles from healthy and COVID-19 BALF, we observed that expression of our Calu-3 gene hits is the most highly modulated upon SARS-CoV-2 infection in the epithelial cell fraction compared to gene hits from other screens **(Figure 3E)**. Taken together, we note strong cell type-specific host factor requirements, with expression of TMPRSS2 playing a major differentiating role. Additionally, screen hits derived from Calu-3 cells were more highly enriched within physiologically relevant human lung epithelial cells.

### Analysis of host restriction factors against SARS-CoV-2

We next sought to systematically investigate the antiviral pathways identified by our GOF screen that are leveraged by the host cell to restrict SARS-CoV-2-mediated cell death. Pathway and protein interaction network enrichment analysis highlighted inflammatory signaling, GPCR signaling, transcriptional regulation, and mucin glycosylation in addition to cell cycle regulation (**Figure 4A and 4B**).

**Figure 4:**
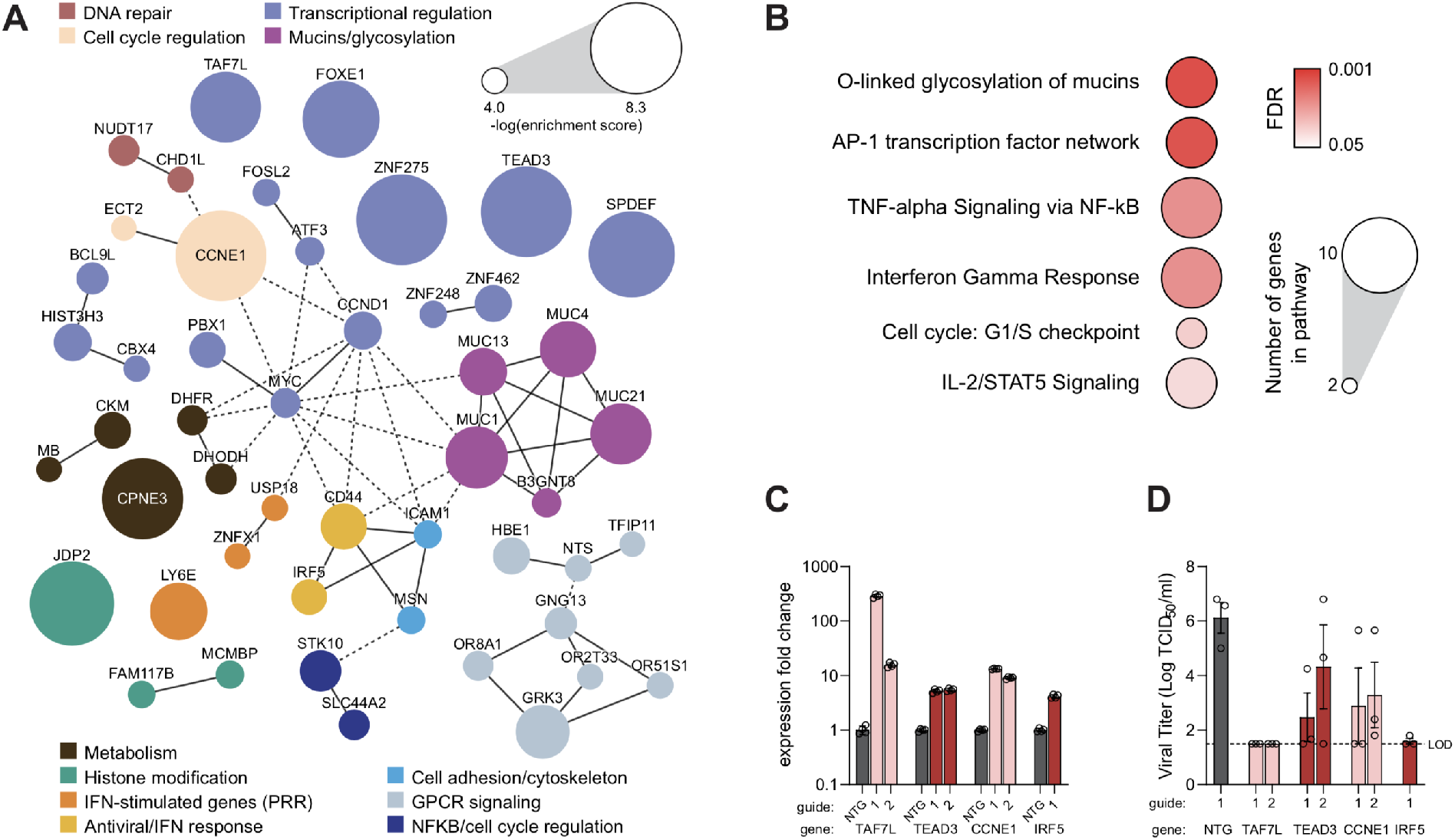
Host restriction factors and pathways of SARS-CoV-2 in lung cells revealed by gain-of-function screening. **A.** Protein-protein interaction network for top 100 enriched hits identified in the CRISPR GOF screen based on STRING analysis. Solid lines between genes indicate direct interaction, dashed lines indicate indirect connections. Nodes are color-coded by functional groups and scaled according to screen enrichment RRA score. **B.** Pathway analysis of top 100 enriched hits indicates significantly overrepresented pathways with putative antiviral roles. Circle size indicates the number of genes within each pathway, color indicates FDR of pathway enrichment. **C.** qRT-PCR analysis of dCas9 transcriptional activation for four screen hits relative to non-targeting controls, n=4. **D.** Individual TCID_50_ validation of the effect of transcriptional upregulation of four putative antiviral hits on SARS-CoV-2 viral titer. Error bars denote mean ± SEM, n=3. Dotted line indicates the limit of detection (LOD) of the assay. NTG = non-targeting guide control.

For secondary validation, we selected enriched GOF hits across these pathways including CCNE1, which regulates the G1-S checkpoint, diverse transcriptional regulators (TEAD3, SPDEF, TAF7L, ZNF248, MRGBP), host helicases (DDX28), ion channels (TMEM206, SLC44A2), and membrane-binding proteins (CPNE3). TCID_50_ assays of these individual GOF Calu-3 cell lines infected with SARS-CoV-2 demonstrated reduced viral titers compared to NTG controls, confirming an antiviral role of these hits (**Figure 4C and 4D, Figure S2A and S2B**).

The antiviral effect of upregulating CCNE1, which drives cells into S phase, is consistent with the proviral effect of activating CDKN2B, a cell cycle inhibitor. Coronavirus regulation of the cell cycle by regulating interactions between cyclins and CDKs has been previously shown for SARS-CoV-1, murine hepatitis virus (MHV), porcine epidemic diarrhea virus (PEDV), and infectious bronchitis virus (IBV), and are a consistent strategy for rerouting host resources towards the coronavirus lifecycle (Su et al., 2020). TEAD3, a transcription factor involved in TGFβ and Hippo signaling, may also regulate cell proliferation in ways that are detrimental to the virus. Another highly enriched transcription factor, SPDEF, is critical for differentiation of goblet cells, a specialized cell type involved in mucus secretion (Chen et al., 2009).

Several of our other GOF enriched hits have been shown to interact with SARS-CoV-2 viral proteins. The DDX28 helicase interacts with the SARS-CoV-2 N protein and was previously reported to be a protective factor in an RNA interference screen for West Nile virus host factors (Chen et al., 2021; Krishnan et al., 2008). CPNE3, a phospholipid-binding protein, has been implicated as an interaction partner with SARS-CoV-1 nsp1 and viral RNA (Cornillez-Ty et al., 2009; Flynn et al., 2020). Transcriptional activation of the transcription factor IRF5 also dramatically reduced viral titer, consistent with the well-established antiviral role of interferon signaling (**Figure 4C and 4D**). Other interferon-stimulated genes (ISGs) identified in the GOF screen were also predicted to result in an antiviral state, including the lymphocyte antigen 6 family member LY6E previously reported to antagonize SARS-CoV-2 entry, ubiquitin specific peptidase (USP18), and the putative mitochondrial-localized dsRNA-sensor ZNFX1 (Honke et al., 2016; Pfaender et al., 2020; Wang et al., 2019).

Finally, we identified mucin glycosylation as a prominent pathway of SARS-CoV-2 host factor restriction **(Figure 4A)**. Mucins comprise a family of high-molecular weight, heavily O-glycosylated glycoproteins and are the primary constituent of mucus lining the epithelial tract of the lungs and gut (Lillehoj et al., 2013). We observed that membrane-tethered mucins (MUC1, MUC4, MUC13, MUC21) form a large, interconnected network in our GOF-enriched protein network analysis along with the acetylglucosaminyltransferase B3GNT6 and cell surface proteins CD44 and ICAM1 **(Figure 4A)**. We decided to focus further validation on transmembrane mucins due to their lung localization, significance of enrichment as an antiviral pathway, and lack of previous studies in the context of SARS-CoV-2 infection.

### Mechanistic insight into the modulation of SARS-CoV-2 cell-entry by mucins

Mucins are ubiquitous proteins encountered by all respiratory and enteric pathogens interacting with the mucosal epithelium. We hypothesized that the mucins identified in our screen are restrictive barriers to SARS-CoV-2 infection of lung epithelial cells. To confirm that upregulation of the membrane-tethered mucins is sufficient to reduce SARS-CoV-2 infection compared to an NTG control, we generated individual GOF cell lines for MUC1, MUC4, MUC13, and MUC21 – unique mucins of varying length and composition – in Calu-3 cells. We also produced a GOF line for the transmembrane glycoprotein CD44, which can be variably spliced to contain a mucin-like domain and interacts with MUC1 (Bennett et al., 1995; Hasegawa et al., 2016). qPCR of the validation cell lines confirmed significant upregulation of target gene transcription by both tested guides in each case **(Figure 5A)**. As predicted, each of these cell lines also exhibited reduced SARS-CoV-2 viral titers compared to NTG controls **(Figure 5B)**. Although this experiment indicates that overexpression of membrane-tethered mucins restricts SARS-CoV-2 infection, it does not reveal if endogenous levels of membrane-tethered mucins modulate SARS-CoV-2 infection. To test this, we treated cells with secreted protease of C1 esterase inhibitor (StcE), a bacterial protease with a distinct peptide- and glycan-based cleavage motif that enables selective cleavage of mucins (Malaker et al., 2019). An enzymatically inactive form of StcE conjugated to Alexa Fluor 647 was used to confirm successful depletion of StcE binding sites **(Figure 5C).** We then performed multi-step growth curves of SARS-CoV-2 in StcE-treated and untreated Calu-3s, to compare SARS-CoV-2 infection kinetics in cells with and without endogenous mucins. We found that SARS-CoV-2 had a growth advantage at 24 hours post infection in StcE-treated cells, compared to untreated control cells, indicating that Calu-3 cells with decreased endogenous mucins were more permissive to viral infection **(Figure 5D)**. Taken together, these data indicate that membrane-tethered mucins restrict SARS-CoV-2 infection *in vitro*.

**Figure 5:**
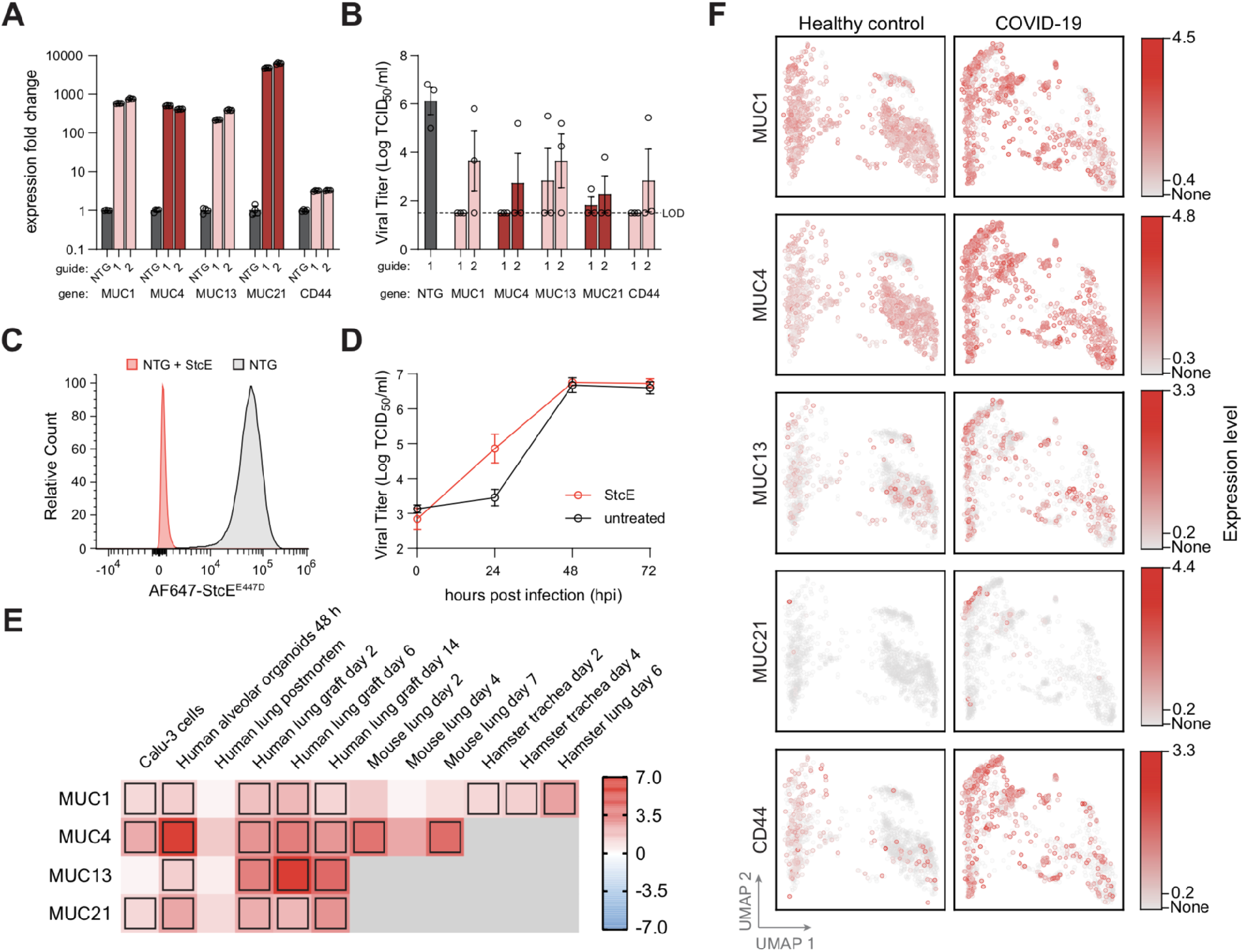
Membrane-tethered mucins are SARS-CoV-2 host restriction factors and upregulated in response to infection. **A.** qRT-PCR analysis of Calu-3 cells overexpressing mucin pathway genes. Fold-change in expression is compared relative to cells transduced with non-targeting guide RNAs (NTG). Error bars denote mean ± SEM, n=3. **B**. Viral titer in GOF cell lines infected with SARS-CoV-2 at MOI 0.05 for 48 hours, as measured by TCID_50_ assay. Two separate sgRNAs were tested per gene. Error bars denote mean ± SEM, n=3. Dotted line indicates the limit of detection (LOD) of the assay. **C.** Flow cytometry data of NTG-transduced Calu-3 cells treated with StcE mucin-selective protease (mucinase). Cells are visualized using a pan-mucin stain conjugated to AF647 dye. **D.** Multi-step growth curves of StcE-digested and untreated Calu-3 cells infected with SARS-CoV-2, as measured by TCID_50_ assay. **E.** Heat map representing differential mucin expression levels in cell models, animal models, and human lung tissue following infection with SARS-CoV-2. Boxes indicate significant differential expression at FDR < 0.05. Color scale indicates log2 fold-change of transcript expression levels **F.** UMAP plots of scRNA-seq data for antiviral host factor expression in lung epithelial cells isolated from BALF of COVID-19 patients and healthy individuals from Liao et al., 2020. Cells are colored based on relative expression levels for each gene. All genes shown exhibit differential expression with FDR < 0.001.

Given their confirmed antiviral role in SARS-CoV-2 infection, we hypothesized that membrane-tethered mucins may be actively regulated in response to SARS-CoV-2 infection. We assessed mucin gene expression in RNA-seq datasets of lung tissues. Across six different model systems in addition to post-mortem human lung tissue, we observe consistent upregulation of mucins from two days up to two weeks post-infection (**Figure 5E**) (Blanco-Melo et al., 2020; Hoagland et al., 2021; Katsura et al., 2020; Muus et al., 2021; Winkler et al., 2020). While hamster and mouse models only exhibited significant upregulation of an individual mucin (MUC1 and MUC4, respectively), human alveolar organoids *in vitro* and human lung grafts *in vivo* exhibited transcriptional activation of all four membrane-tethered mucins. We further analyzed single cell RNA-seq data of COVID-19 patient BALF for active transcriptional regulation of validated GOF hits in response to infection. The epithelial cell fraction of COVID-19 BALF revealed significant upregulation of all 4 transmembrane mucins in addition to CD44 and TEAD3 (**Figure 5F**). Taken together, our data indicates that mucin upregulation is a broad antiviral host response across multiple species.

We next sought to identify the stage of the viral life cycle that is affected by mucin upregulation. As all four mucins enriched in our GOF screen are expressed at the host cell surface, we hypothesized that they reduce SARS-CoV-2 entry. To test this, we utilized vesicular stomatitis virus (VSV) encoding GFP and pseudotyped with SARS-CoV-2 spike protein (VSV-S) to measure the effect of mucins on Spike-mediated viral entry (**Figure 6A**). Live cell imaging demonstrated that overexpression of all tested membrane-tethered mucins, including CD44, inhibited VSV-S infection relative to NTG control, in agreement with our viral titer validation data **(Figure 6B, Figure 5B)**.

**Figure 6:**
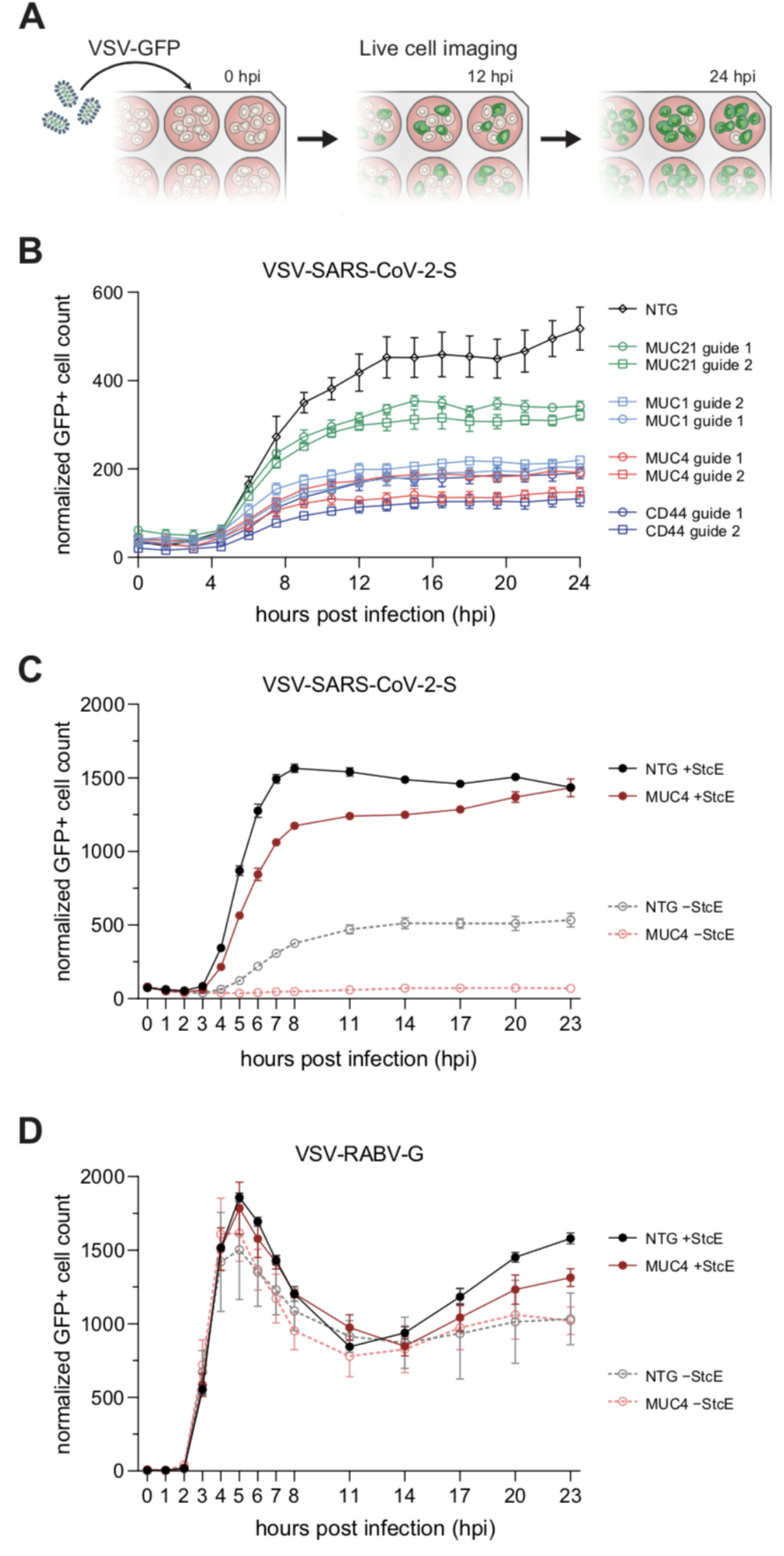
Membrane-tethered mucins restrict SARS-CoV-2 Spike-mediated entry. **A.** Schematic of the pseudotype entry assay using a SARS-CoV-2 Spike-pseudotyped vesicular stomatitis virus (VSV-S) encoding GFP. Following infection, cells are imaged regularly over 24 hours to track viral infection by quantifying GFP^+^ cell counts. **B.** Timecourse of VSV-S infection of NTG, MUC1, MUC4, MUC21, and CD44-overexpressing GOF cell lines. NTG, non-targeting guide. Error bars denote mean ± SEM, n=3. **C.** Timecourse of VSV-S infection of NTG and MUC4 GOF cells pre-treated with StcE mucinase prior to VSV-S infection. Cells exhibited increased spike-mediated viral entry into cells when compared to untreated cells. Error bars denote mean ± SEM, n=3. **D.** Timecourse of VSV-RABV-G entry in NTG and MUC4 GOF cells pre-treated with StcE mucinase prior to pseudotype virus infection. VSV-RABV-G, VSV particles pseudotyped with rabies virus glycoprotein. Error bars denote mean ± SEM, n=3.

To confirm that the decreased VSV-S infection observed is mediated by extracellular mucin coats, we selected MUC4 for further study due to its strong phenotype in our TCID_50_ assay. We treated MUC4 GOF and NTG cell lines with StcE to determine if removal of overexpressed MUC4 and endogenous mucins modulated VSV-S infection. Interestingly, StcE digestion of both cell lines led to a dramatic increase in VSV-S infection compared to untreated cells, indicating that mucin removal renders cells more permissive to VSV-S infection **(Figure 6C)**. In addition, the lower VSV-S infection in MUC4 GOF cells, relative to the NTG controls, were partially reversed in StcE treatment conditions **(Figure 6C)**. These data further indicate that removal of endogenous mucins from Calu-3 cells, in addition to overexpressed mucins, markedly increases VSV-S infection. Calu-3 cells endogenously co-express multiple membrane anchored mucins including both MUC1 and MUC4. Further, no mucins emerged from our LOF screen, indicating that distinct transmembrane mucins may compensate for each other. This underscores the unique advantage of GOF screens in elucidating antiviral host-pathogen interactions. To confirm whether mucin inhibition of VSV-S infection was specific to the SARS-CoV-2 S glycoprotein, we also tested VSV pseudotyped with the rabies virus glycoprotein (VSV-RABV-G). In contrast to VSV-S, we did not observe an inhibitory effect on infection by VSV-RABV-G of either endogenously expressed mucins or upregulated MUC4 in Calu-3 cells, confirming that VSV-S infection inhibition in mucin GOF Calu-3s was specific to S-mediated entry **(Figure 6D).** Taken together, our data indicate that membrane-tethered mucins restrict SARS-CoV-2 entry both when present at endogenous levels as well as when overexpressed and suggest that they may modulate SARS-CoV-2 infection in COVID-19 patients.

## DISCUSSION

Here, we conducted genome-scale CRISPR loss- and gain-of-function screens in human lung epithelial cells, identifying numerous host factors that promote or restrict SARS-CoV-2-mediated cytopathic effects. In addition to reporting the first GOF screen with SARS-CoV-2, we present the first LOF screen in human lung cell types that have not been engineered with entry factors to improve the efficiency of viral infection. Hits identified in our Calu-3 knockout screen were found to be highly enriched in ciliated ACE2^+^/TMPRSS2^+^ lung epithelial cells from a meta-analysis of single-cell RNA sequencing studies from over 200 individuals; these hits are largely divergent from hits identified in other recent LOF screens. This host factor catalog is thus likely to include hits that are unique to the TMPRSS2-dependent cell entry route of SARS-CoV-2 infection in human lung epithelial cells (**Tables S1-S3**). In contrast, other CRISPR LOF screens have identified genes critical for a CTSL-dependent entry route, including factors involved in vascular trafficking and acidification of vacuoles (Bailey and Diamond, 2021; Daniloski et al., 2021; Flynn et al., 2020; Schneider et al., 2021; Wang et al., 2021; Wei et al., 2021; Zhu et al., 2021).

A strength of bi-directional CRISPR screens is that genes and pathways can be interrogated in both a GOF- and LOF-manner simultaneously, possessing potential to capture the complexity of host-pathogen interactions. The emergence of inversely complementary hits from GOF and LOF screens (*e.g*. key pathway activators and inhibitors emerging from opposite screen directions) increases confidence in the importance of a given pathway for SARS-CoV-2 infection. Our screens identified several such pathways including cell cycle, NF-κB signaling, and epigenetic modifiers.

While many DNA viruses such as papillomaviruses, adenoviruses, and polyomaviruses actively promote the cell cycle to increase availability of DNA replication machinery (Bagga and Bouchard, 2014), the opposite trend is generally seen for RNA viruses. Several members of the *Coronaviridae* family including SARS-CoV-1 and murine hepatitis virus (MHV) employ diverse strategies for cell cycle arrest, which is hypothesized to benefit the virus via redistribution of cellular resources, avoidance of apoptosis-triggering checkpoints, or maintenance of organelle architecture required for viral replication (Dyer et al., 2008; Fan et al., 2018; Su et al., 2020; Yuan et al., 2006). SARS-CoV-2 likely exploits similar approaches, although the specific mechanisms are still emerging. Previous work has demonstrated SARS-CoV-2 genomic RNA directly interacts with the cell cycle promoter CDK2, which also exhibits decreased phosphorylation during infection (Bouhaddou et al., 2020). We observed that upregulation of factors promoting the cell cycle such as cyclin E1 and cyclin D1 restrict SARS-CoV-2 infection in Calu-3 cells, while overexpression of cell cycle inhibitors like CDKN1A and CDKN2B enhance SARS-CoV-2 infection. Although some screen hits may be enriched in pooled survival screens due to their effect on cell proliferation independent of viral infection, we validated CCNE1 and CDKN2B in assays directly measuring viral replication. Supporting this, cell cycle-modulating compounds such as dinaciclib and AZD5438 antagonize SARS-CoV-2 replication (Biering et al., 2020; Bouhaddou et al., 2020).

Taken together, a growing body of work indicates cell cycle arrest is critical for optimal SARS-CoV-2 infection (Cawood et al., 2007; Mizutani et al., 2006; Yuan et al., 2006, 2007).

NF-κB-mediated signaling also emerged from bidirectional functional screening. Some factors, such as NF-κB-induced genes CD44 and ICAM1, emerged as putatively antiviral, based on enrichment in the GOF screen. However, the majority of the NF-κB related hits appeared as proviral, with CHUK/IKK-α and RIPK4 enriched in the LOF screen and NFκB pathway activators IL-17C and its signaling adapter protein ACT1 (TRAF31P2) depleted in the GOF screen. Human coronavirus 229E selectively modulates certain components of the NF-κB signaling pathway for its replicative benefit in an IKK complex-dependent manner (Poppe et al., 2017). A proviral role of the NF-κB pathway has also been reported for influenza A virus (IAV), as NF-κB inhibitors as well as genetic perturbation were demonstrated to restrict IAV infection *in vitro* and *in vivo* (Ehrhardt et al., 2013; Kumar et al., 2008; Pinto et al., 2011; Schmitz et al., 2014). A similar mechanism for a proviral role of proinflammatory cytokines and NF-κB-signaling for SARS-CoV-2 would be plausible. Alternatively, a more indirect role via modulation of cell proliferation or cell death pathways would also be possible.

One of the most prominently enriched proviral clusters from our LOF screens was a set of clathrin adaptor proteins from the AP1 complex, which are involved in vesicle formation and intracellular trafficking. Several AP1 adaptor proteins have also been identified as interactors of SARS-CoV-2 M protein, including one of our top validated hits, AP1G1 (Chen et al., 2021). AP1 adaptors could either serve a direct role in the viral life cycle by affecting intracellular viral trafficking and egress or be manipulated by the SARS-CoV-2 M protein to reroute host proteins and facilitate viral infection. A similar mechanism is used by HIV-1, which hijacks the AP1 complex to retain the viral restriction factor BST2 within endosomes and drive its lysosomal degradation (Jia et al., 2014). Depletion of AP1 components can also alter cell surface expression of membrane proteins (Castillon et al., 2018; Takahashi et al., 2011). As a result, entry factors like ACE2 and TMPRSS2 may be mislocalized in AP1 LOF cells, resulting in decreased viral entry. Disrupting the AP1 complex *in vivo* has also been demonstrated to result in altered epithelial cell-cell junctions and cell polarity, resulting in a hyperproliferative state (Bonifacino, 2014; Boyden et al., 2019; Hase et al., 2013). Given our data suggesting a critical role for cell proliferation regulation and cell-cell junction proteins like E-cadherin (CDH1) in SARS-CoV-2 infection, perturbation of these pathways in AP1 LOF cells may also explain their resistance to SARS-CoV-2 infection (Bonifacino, 2014; Boyden et al., 2019; Castillon et al., 2018; Hase et al., 2013; Takahashi et al., 2011).

Another central class of genes identified in our LOF screen were components involved with cytoskeletal rearrangement (ROCK1) and cell-cell junctions (E-cadherin). Viruses commonly coopt actin or microtubule components and motor proteins to shuttle viral components around the cell and enable efficient replication (Arons et al., 2020; Taylor et al., 2011; Wen et al., 2020). Beyond intracellular shuttling, some viruses hijack host cytoskeletal components as well as cellcell protein complexes for cell-cell spread (Bergelson, 2009; Bouhaddou et al., 2020; Gordon et al., 2020; Mothes et al., 2010). An example of this for SARS-CoV-2 is the observation that SARS-CoV-2 nsp7 and N proteins mediate cytoskeletal rearrangement by signaling through RhoA and CK2, forming filopodial protrusions from infected cells that have been hypothesized to serve as portals of cell-cell spread (Bergelson, 2009; Bouhaddou et al., 2020; Gordon et al., 2020).

Furthermore, as ACE2 has been reported to be preferentially localized to the cilia of upper and lower respiratory tract epithelial cells, loss of cytoskeletal components could also affect proper cell surface expression of ACE2 and potentially other SARS-CoV-2 entry factors, subsequently decreasing viral entry (Lee et al., 2020). Further, because cell division involves a tight coordination between the cytoskeleton, cell-cell junctions, and nuclear components, some of the identified cell junction and cytoskeleton components may also regulate the cell proliferative state, potentially connecting these two pathways.

Our GOF screen identified several G-protein coupled receptors involved in smell and taste signaling as SARS-CoV-2 restriction factors. A key protein interaction network enriched in the GOF screen include the olfactory receptors OR8A1, OR2T33, and OR51S1, the G protein gamma subunit GNG13 that plays a key role in taste transduction, and the GPCR kinase GRK3. Respiratory epithelium and neuroepithelium of the human nasal cavity robustly express ACE2 and TMPRSS2, and transcripts encoding olfactory receptors and GNG13 are downregulated in human and hamster olfactory epithelium following SARS-CoV-2 infection (Brann et al., 2020; Fodoulian et al., 2020; Zazhytska et al., 2021). The specific mechanism of an antiviral role for olfactory receptors, and their potential connections to COVID-19 symptoms such as anosmia, will warrant further investigation. However, many sensory receptors are expressed in diverse tissues and have functions beyond their canonical role in olfactory epithelium sensory neurons. Taste receptors in airway epithelia can play a direct protective role, sensing inhalation of noxious chemicals as well as metabolic products of pathogens. By activating calcium signaling and other second messenger pathways, they then increase cilia beat frequency to drive mucociliary clearance (Lee et al., 2012; Shah et al., 2009). Furthermore, whole exome studies of infants infected with respiratory syncytial virus (RSV) identified genetic variants in OR8U1 and OR8U8, suggesting that loss of some olfactory receptors could be involved in increased host susceptibility to RSV infection (Salas et al., 2017). Mutations in our antiviral screen hit GRK3 have also been found to be associated with immunodeficiency disorders (Balabanian et al., 2008). Because we did not identify olfactory or gustatory components in the LOF screen, which may not be expressed in lung epithelial cells such as Calu-3 lines, GOF screens highlight the potential to find novel functional pathways via ectopic overexpression.

We also identified membrane-tethered mucins forming a prominent interaction cluster of SARS-CoV-2 restriction factors. Mucins are a family of densely O-glycosylated proteins that are the primary constituent of mucus lining epithelial cell barriers in the lungs and gut (Wagner et al., 2018), protecting the respiratory tract from environmental insults such as microbial infection (Button et al., 2012; Kesimer et al., 2013). Mucins are highly expressed by epithelial cells (Ma et al., 2018) and are separated into two major classes. Secreted, gel-forming mucins form the mucus layer (such as MUC5AC, MUC5B), while transmembrane mucins (such as MUC1, MUC4, MUC13, MUC21) are anchored to the apical side of epithelial cells (Carson, 2008; Hattrup and Gendler, 2008). Viral infections, including by SARS-CoV-2, have been shown to upregulate mucin expression in primary lung epithelial cells in a type I IFN-dependent manner, suggesting potential protective roles for mucins (Iverson et al., 2021; Liu et al., 2020; Lu et al., 2021). Our investigation demonstrates that membrane-tethered mucins serve as SARS-CoV-2 restriction factors in Calu-3 cells, inhibiting infection at the stage of viral entry **(Figure 5-6)**. This is supported by quantitative trait locus (QTL) and *in vivo* studies suggesting that membrane-tethered MUC4 could serve a protective role in mice against SARS-CoV-1 and chikungunya virus infection (Plante et al., 2020).

In the context of SARS-CoV-2, mucin studies have largely focused on their potentially detrimental functions during the later stages of COVID-19. Overproduction and accumulation of secreted mucins in the lungs can lead to airway obstruction, thereby reducing airflow, exacerbating lung disease, and potentially promoting ARDS (He et al., 2020; Nakashima et al., 2008; Vestbo, 2002). It has therefore been proposed to antagonize mucin expression as a therapeutic strategy (Guan et al., 2020; Kost-Alimova et al., 2020; Liu et al., 2020), and compounds that inhibit MUC1 are now being tested in clinical trials for hospitalized adults with COVID-19 (Strich, 2020; Tabassum et al., 2020).

Given our findings of a protective role for membrane-anchored mucins against SARS-CoV-2, the development of therapeutics broadly antagonizing mucin expression should be pursued with caution. Structural variant studies of COVID-19 cases identified MUC4 loss-of-function to be associated with increased disease severity (Reay et al., 2021; Sahajpal et al., 2021). A more selective strategy promoting membrane-tethered mucins or reducing the secretion of gel-forming mucins may be more beneficial. In line with this, the sole secreted mucin (MUC5AC) identified in our screens was depleted in the GOF study, suggesting a proviral role. Taken together, the levels of distinct mucins in the lung of infected patients may determine the balance between the protective benefit of these surface glycoproteins and the pathological detriment in gas exchange.

Future work is required to determine the exact mechanism of membrane-anchored mucins in restricting SARS-CoV-2 entry. In addition to direct steric hindrance of S glycoprotein receptor binding, the glycan composition of some mucins could potentially enhance binding of SARS-CoV-2 virions to the cell surface, given that SARS-CoV-2 interactions with membrane glycosaminoglycans (GAGs) are important for viral entry (Clausen et al., 2020; Nguyen et al., 2021; Zhang et al., 2020). In contrast, Influenza A virus (IAV) interacts with sialic acid residues on host mucins, which can actively trap viral particles and inhibit viral entry (Cohen et al., 2013; Delaveris et al., 2020; Ehre et al., 2012; Iverson et al., 2021; McAuley et al., 2017). Defining the molecular interactions between SARS-CoV-2 virions and mucins, and their consequences during SARS-CoV-2 infection *in vivo*, will clarify the potentially bimodal effect of mucins on COVID-19 severity.

In summary, we present a systematic functional catalog of highly interconnected host factors and pathways that mediate SARS-CoV-2 infection in human lung cells including cell cycle modulation, intercellular junction dynamics, clathrin-mediated trafficking, inflammatory responses, and O-linked glycosylation of mucins. The combination of bidirectional loss- and gain-of function genome-scale screens enables the assignation of proviral or antiviral roles for individual genes in these complex pathways. By exploiting canonical ACE2 and TMPRSS2-mediated viral entry into lung epithelial cells with endogenous expression of these entry factors, we were able to identify unique host factor dependencies with robust enrichment in single cell RNA sequencing datasets of healthy and infected human lung epithelium. We then probed the roles of membrane-tethered mucins in restricting SARS-CoV-2 infection and demonstrated that restriction occurs at the stage of viral entry. By dissecting the interactions between SARS-CoV-2 and lung epithelial cells, these host factor screens provide a starting point for host-directed therapeutic intervention.

## Acknowledgements

We thank Dr. Richard C. Boucher (University of North Carolina) for critical insight and discussion on mucin biology. We thank Dr. Britt Glaunsinger (UC Berkeley), Dr. Patrick S. Mitchell (UC Berkeley/University of Washington), and Dr. Michael S. Diamond (Washington University School of Medicine) for helpful discussion and critical reading of this manuscript. We thank Dr. Ella Hartenian (UC Berkeley), Dr. David Morgens (UC Berkeley), and Dr. Wei Li (Children’s National Hospital) for helpful discussion. We thank the UC Berkeley Cell culture facility for providing Calu-3 cell lines and Dr. Melina Fan (Addgene) for assisting with critical plasmid and lentivirus resources. The USA-WA1/2020 SARS-CoV-2 clinical isolate, NR-52281 was obtained from BEI resources, NIAID, NIH and deposited by the Centers of Disease Control and Prevention. S.B.B. is an Open Philanthropy Awardee of the Life Sciences Research Foundation. E.V.D. is supported by NSF Graduate Research Fellowship DGE-1752814. J.C.S. is supported by an NIH/NCI F32 Postdoctoral Fellowship 1F32CA250324-01. D.J.S. is supported by an NSF Graduate Research Fellowship. This work was supported, in part, by National Cancer Institute Grant R01CA200423 (to C.R.B.). This work was supported in part by N.I.H. R01 AI140186 and Burroughs Wellcome Fund (Investigators in the Pathogenesis of Infectious Disease) to J.E.C.. S.K. is a Howard Hughes Hanna Gray fellow and Chan Zuckerberg Biohub investigator. This work was supported by Fast Grants (to P.D.H. and S.A.St) and NIH (P.D.H. DP5OD021369) funding.

## Author Contributions

P.D.H., S.B.B., and E.H. conceived the study. S.B.B., S.K., S.A.S., E.W., and P.D.H. led experimental design. S.A.S., E.W., and S.B.B performed BSL2 and molecular biology experiments. S.B.B., X.N., E.V.D., and L.H.Y. performed BSL3 experiments. J.R.Z. performed pseudotype entry assays with input from J.E.C.. S.K., V.S., C.C., A.B., and P.D.H. performed computational analysis. J.C.S, D.J.S., and C.R.B. supported mucin validation experiments. D. M.F. and S.A.St. supported BSL3 experiments. P.D.H. provided overall supervision along with E. H. and S.K.. S.B.B., S.K., and P.D.H. wrote the manuscript with help from all authors.

---

**Supplemental Figure 1:**
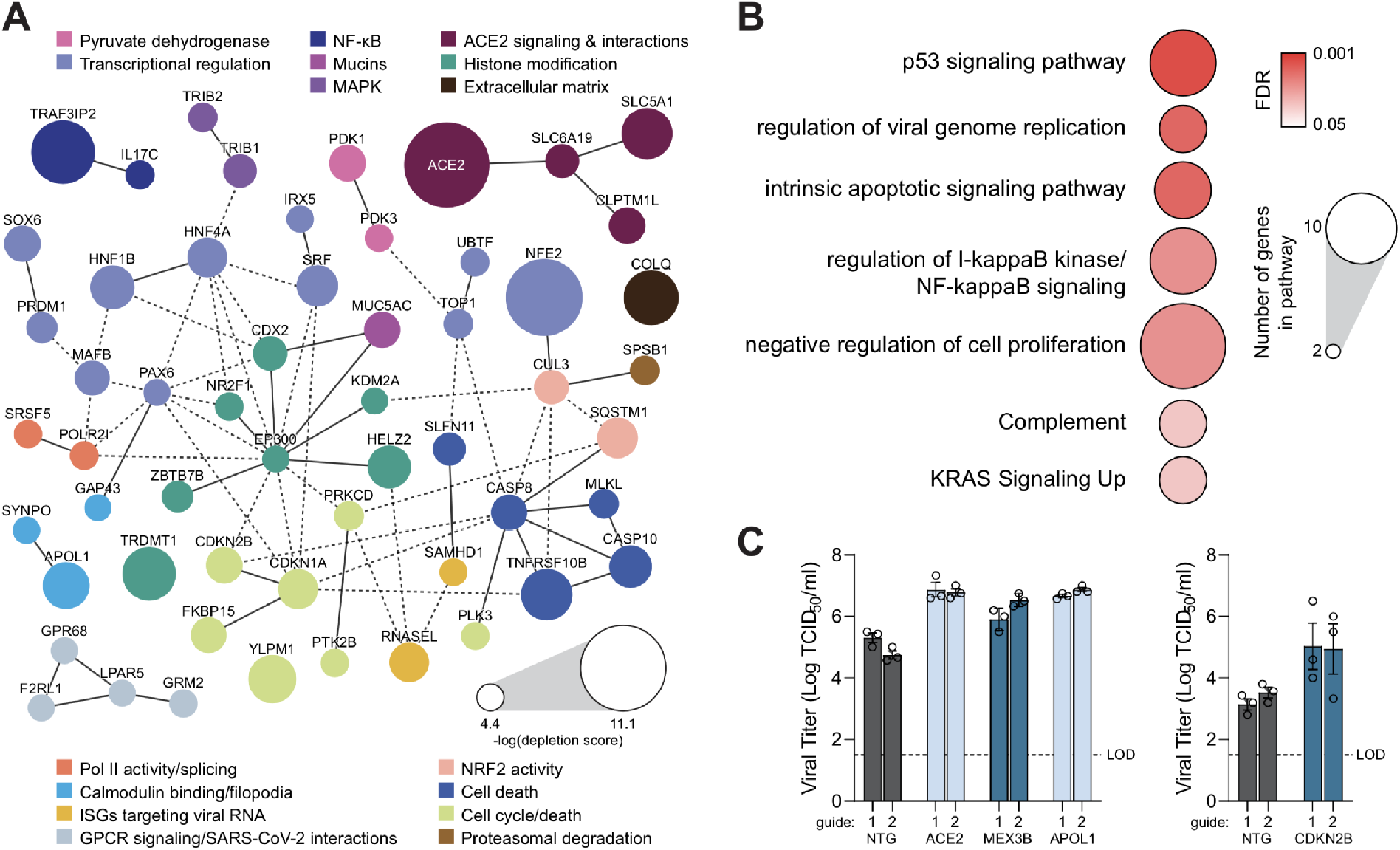
SARS-CoV-2 proviral signatures revealed by depleted genes in the GOF screen. **A.** Pathway analysis of top 100 depleted hits in the GOF screen indicates significantly overrepresented pathways with putative antiviral roles. Circle size indicates the number of genes within each pathway, color indicates FDR of pathway enrichment. **B.** Protein-protein interaction network for top 100 depleted hits identified in the CRISPR GOF screen based on STRING analysis. Solid lines between genes indicate direct interaction, dashed lines indicate indirect connections. Nodes are color-coded by functional groups and scaled according to screen enrichment RRA score. **C.** Individual guide TCID_50_ validation of the effect of transcriptional upregulation of four putative proviral hits on SARS-CoV-2 viral titer in Calu-3 cells infected at an MOI of 0.05 for 48 hours. Each gene was targeted with two sgRNAs. Error bars denote mean ± SEM, n=3. Dotted line indicates the limit of detection (LOD) of the assay. NTG = non-targeting guide control.

**Supplemental Figure 2:**
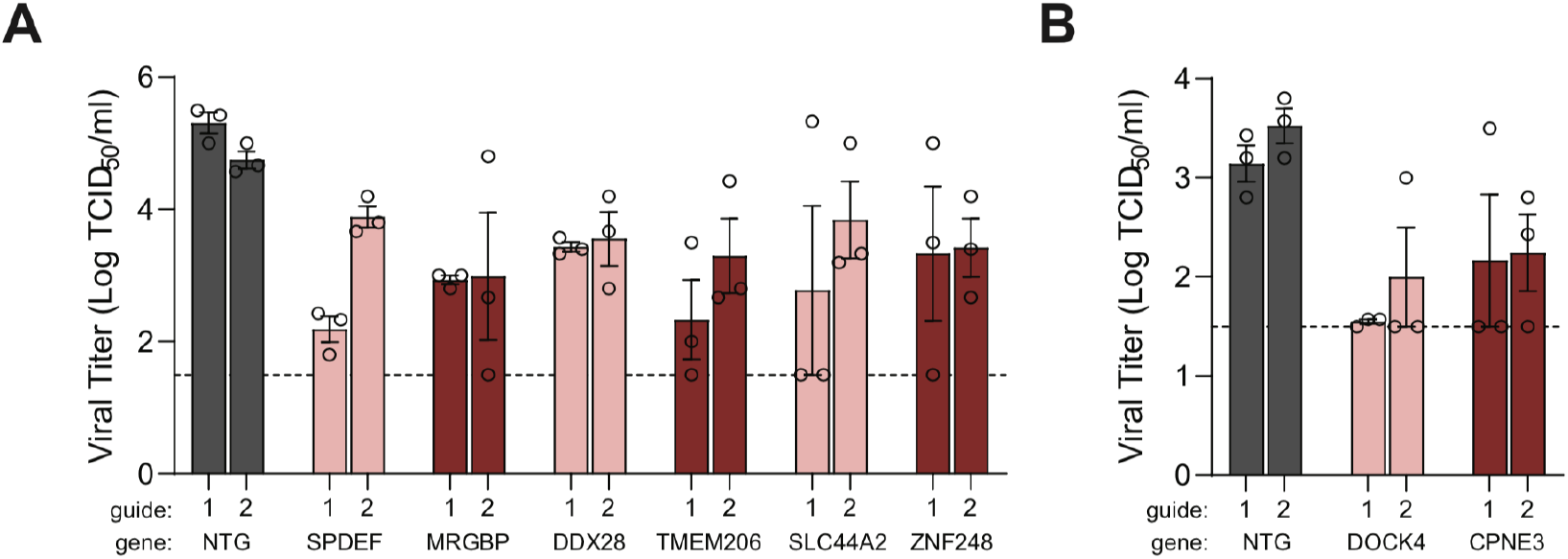
Validation of additional SARS-CoV-2 restriction factors enriched in the GOF screen. **A.** Viral titer in GOF cell lines infected with SARS-CoV-2 at an MOI of 0.05 for 48 hours, and measured by the TCID_50_ assay. Two separate sgRNAs were tested per gene hit. Error bars denote mean ± SEM, n=3. Dotted line indicates the limit of detection (LOD) of the assay. **B.** A separate round of TCID_50_ assays were conducted as in the previous panel. Error bars denote mean ± SEM, n=3.

**Supplementary Table 1: LOF enriched screen analysis**

**Supplementary Table 2: GOF depleted screen analysis**

**Supplementary Table 3: GOF enriched screen analysis**

**Supplementary Table 4: Sequences used in this study**

## METHODS

### Cell culture

The human lung epithelial cell line Calu-3 (UC Berkeley Cell Culture Facility) was maintained RPMI media supplemented with Glutamax (Thermo Fisher), 10% FBS (HyClone), and 100 μg/ml penicillin and streptomycin (Gibco) [R10]. Cells were grown in T225 flasks (Thermo Fisher) and regularly passaged at 80-90% confluence with 10x concentrated Tryple Express (Thermo Fisher). After lentiviral transductions with Cas9 and dCas9-VP64 constructs, cells were cultured in R10 media additionally supplemented with 16 ng/mL of hepatocyte growth factor (HGF, Stem Cell Technologies) to preserve viability and support robust growth (Datta et al., 2017).

### Guide RNA library amplification

Endura electrocompetent cells (Lucigen) were electroporated with the genome-scale CRISPR KO (Brunello) and CRISPR activator (Calabrese) libraries, a gift from John Doench (Addgene). After recovery, cells were grown on LB agar bioassay plates supplemented with 100ug/mL carbenicillin overnight at 37C and harvested at a coverage of >1000 colonies per guide. Library plasmid DNA was extracted using a Nucleobond EF maxiprep kit (Macherey Nagel) followed by Novaseq sequencing to verify guide RNA representation.

### Lentiviral production

Low-passage HEK 293FT cells were grown in DMEM supplemented with 10% FBS (D10 media) and passaged using TrypLE (Gibco). For viral plasmid transfection, Polyethylenimine ‘Max’ (PEI, linear, MW 40,000, VWR) at a concentration of 1mg/mL and pH 7.1 was used. PEI (0.71ug PEI/cm^2^ cell culture area) was mixed with 1 mL of DMEM. pMD2.G, PAX and library plasmid (0.178ug/cm^2^, at a 0.25 (pMD2.G): 0.5 (psPAX): 1(Library plasmid) ratio were thoroughly mixed with 1 mL DMEM. Following 10 min of incubation at room temperature, the DNA and PEI mixtures were combined, mixed and incubated for 30 min at room temperature. 150,000 cells/cm^2^ of 293FT cells in suspension were mixed with the PEI-DNA DMEM mixture and allowed to incubate for 5 minutes at room temperature and then plated in D10 for 48 hours. Lentiviral supernatant media was harvested, filtered through a 0.45um filter and frozen at −80°C prior to titration on Calu-3 cells.

### SARS-CoV-2 stock preparation and infections

The USA-WA1/2020 strain of SARS-CoV-2 was used in this study which was obtained from BEI Resources. The original stock from BEI was passaged through a 0.45 μM syringe filter and then 5 μl was inoculated onto an 80% confluent T175 flasks (Nunc) of Vero-E6 cells to produce our p1 stock. Cytopathic effect was monitored daily and flasks were frozen when cells exhibited ~70% cytopathic effect, usually ~48-72 hpi. Lysates were then thawed, collected, and cell debris was spun down at 3000 x rpm for 20 minutes. The clarified virus containing supernatants were then aliquoted and infectious viral particles were quantified with a TCID50 assay. To produce p2 working SARS-CoV-2 stocks, 5 μl of the p1 stock was inoculated onto 80% confluent T175 flasks of Vero-E6 cells as described above.

### Genome-wide, bidirectional CRISPR screens

Calu-3 cells were transduced with lenti Cas9-Blast (Addgene #52962), or with lenti dCAS-VP64_Blast (Addgene #61425), gifts from Feng Zhang. 24 hours post-transduction, cells were selected with 10ug/mL of blasticidin for 10 days. Cas9 and dCas-VP64 Calu-3 knock-in lines were then transduced with Brunello and Calabrese Set A libraries (Addgene #73179 and #92379) as appropriate, which were gifts from John Doench. Library transductions were conducted to maintain >1000X representation of sgRNAs. The Brunello library contains 76,441 sgRNAs, so >2.7×10^8^ cells were transduced at MOI 0.3. The Calabrese library contains 56,762 sgRNAs, so >2×10^8^ cells were transduced at MOI 0.3. 24 hpi, cells were selected with 1.5 ug/mL for 5 days.

For the LOF screen, the Cas9 knock-in cell line transduced with Brunello sgRNAs were seeded at 3.5×10^7^ cells per T225 flask and allowed to grow to 70% confluency. At this point, half of the cells were harvested for a Day 0 (D0) timepoint to serve as a reference for sgRNA enrichment analysis. The remaining cells were infected with the SARS-CoV-2 USA/WA-1 isolate at an MOI of 0.05. Cells were incubated for 4 days until >70% CPE was apparent. Throughout infection, media was changed every other day with R10+HGF. For the GOF screen, dCas-VP64 knock-in cells transduced with Calabrese sgRNAs were treated similarly except they were infected at an MOI of 0.05 for 5 days until >70% CPE was apparent. At 5 days post infection, surviving cells were uplifted and lysed according to manufacturer instructions using a genomic DNA extraction kit, followed by heat inactivation (Zymo Research).

### Next-generation sequencing

Guide RNA cassettes were amplified from extracted genomic DNA to generate Illumina sequencing libraries. Namely, 3 ug of genomic DNA was added per 50uL PCR reaction using staggered primers to increase base diversity. PCR products were then pooled and purified using QIAquick PCR purification kits (Qiagen). Different libraries were then quantified with Kappaquant to determine relative concentrations of amplified product, then pooled to match concentrations. Pooled samples were then sequenced by Illumina Novaseq. In addition to the uninfected D0 and infected timepoints, the lentiviral plasmid prep was also sequenced to assess any guide distribution skew resulting from transduction.

### Computational analysis of CRISPR screens

A previously published method for CRISPR screen analysis, MAGeCK, (Li, Wei et al. 2014) was used to rank genes based on redundant targeting guides via robust rank aggregation (RRA). Both GOF and LOF screens were performed with 4 replicates. sgRNA enrichment and depletion was assessed in infected vs. D0 uninfected samples for each paired sample using the --paired flag. Each set of top 100 screen hits was determined based on MAGeCK RRA rank and the false discovery rate in a negative control analysis. To investigate the relationships within each of these sets, we performed protein-protein interaction networks functional enrichment analysis with STRING (Szklarczyk, Damian et al. 2019). Any genes that exhibited depletion or enrichment in the top 500 hits between the plasmid library and the uninfected control were removed from STRING analysis as they are likely confounded by affecting cellular growth or survival unrelated to viral infection. Nodes were resized based on MAGeCK scores. Specifically, the radius (*r*) of each node was calculated as a constant factor (*c*) scaled by 1.5 raised to the z-score (*z*) of the corresponding gene’s negative log MAGeCK score: *r* = *c* * 1.5^z^. *z* was calculated from the distribution of the negative log MaGeCK scores in the relevant top 100 gene set.

### Secondary validation with individual guide RNAs

Lentivirus for Cas9 KO was produced in 6-well plates (scaled down from the lentivirus production protocol). After 48 hours, viral supernatant was harvested, centrifuged at 800xg for 5 minutes to remove cell debris, and used to transduce 800,000 Cas9-expressing Calu-3 cells. 24h later, cells were selected with Blasticidin to completion. Lentivirus for CRISPR activators was produced as described above, scaled to T75 flasks. Clarified lentiviral supernatant was concentrated using LentiX Concentrator (Takara Bio). The concentrated aptamer guide RNA virus was used to transduce 5M Calu-3 cells expressing dCas9-VP64 prior to blasticidin selection the following day.

### Calu-3 validation infections and TCID50 assay

2×10^5^ Calu-3 cells were seeded into 24-well plates and infected with SARS-CoV-2 48 hours postseeding at an MOI of 0.05. Viral inoculums were incubated with cells for 30 minutes at 37 C, at which time they were removed and cells were washed once with 1xPBS. Regular media R10+HGF was then replaced and cells were incubated for the times indicated in the figures (24-48 HPI). Plates were then frozen down to lyse cells and then thawed when ready to quantify virus by TCID50. To titer infectious viral particles, plates were thawed and viral lysates was serially diluted, and each dilution was applied to eight wells in 96-well plates containing Vero-E6 cells. Three days later, CPE was counted visually and TCID50/mL was calculated using the dilution factor required to produce CPE in half of the wells (4/8) for a given dilution.

### qPCR validation of overexpressing cell lines

Cells were cultured in 96-well plates until >70% confluency, and cDNA was acquired with direct lysis to reverse transcription protocol as described in (Joung et al., 2017). Taqman qRT-PCR was completed with the following probes: TAF7L (Hs00227589_m1), TEAD3 (Hs00243231_m1), CCNE1 (Hs01026536_m1), IRF5 (Hs00158114_m1), MUC1 (Hs00159357_m1), MUC4 (Hs00366414_m1), MUC13 (Hs00217230_m1), MUC21 (Hs01379324_g1), CD44 (Hs01075861_m1). GAPDH was used as an endogenous control gene.

### Generation of replication competent vesicular stomatitis virus (VSV) pseudovirus

Recombinant VSV expressing eGFP and SARS-CoV-2-S (VSVdG-eGFP-CoV-2-S) was generated as previously described (Wang et al., 2021). In short, VSVdG-eGFP (Addgene, Plasmid #31842) was modified to insert in frame with the deleted VSV-G a codon optimized SARS-CoV-2-S based on the Wuhan-Hu-1 isolate (GenBank:MN908947.3), which was mutated to remove a putative ER retention domain (K1269A and H1271A). The virus was rescued and passaged in Huh7.5.1 cells multiple times until widespread GFP fluorescence and cytopathic effects were observed. Virus was propagated on Vero-E6 cells and titrated on VeroE6 cells overexpressing TMPRSS2. Sequencing revealed additional mutations at the C-terminus (1274STOP) and a partial mutation at A372T (~50%) in the ectodomain. Similar adaptive mutations were found in a previous published VSVdG-CoV-2-S (Dieterle et al., 2020).

Recombinant VSV expressing eGFP and rabies virus G (VSVdG-eGFP-RABV-G) was generated in a similar manner. Rabies virus (RABV) glycoprotein (G) was amplified from a template containing the G sequence from SAD-B19. This gene was assembled into VSV-eGFP-dG (Addgene, Plasmid #31842) in frame with the G coding sequence between MluI and NotI to generate VSV-eGPF-RABV-G. To rescue the VSVdG-RABV-G, 293FT cells (Thermo Fisher Scientific) were co-cultured with VeroE6 cells in a 6-well plate. Cells were transfected with pCAGEN-VSV-N (300ng), pCAGEN-VSV-P (500ng), pCAGEN-VSV-L (200ng), pCAGEN-VSV-G (800ng), pCAGGS-T7 (200ng), and VSV-eGFP-dG-RABV-G (500ng) using JetPrime (Polyplus). Media was changed to DMEM + 2% FBS after 1 day, and cells were observed until widespread GFP was observed by day 7. Rescued virus was plaque purified, propagated, and titrated on Vero-E6 cells. Sequencing confirmed that the sequence matched the SAD-B19 and no mutations were observed.

### Quantifying pseudovirus infection

Cells were plated in clear 96-well plates for infection and 50-80% confluence the next day. Media was removed from cells and pseudovirus was added at an MOI of 0.1 in 200 μl of DMEM+2% FBS. Plates were spun at 900xg at 30°C for 60 minutes. After spinfection, individual wells were imaged over time using an Incucyte System (Sartorius) in a 37°C incubator at 5% CO_2_. Images were taken of each well at 4x magnification, and infection was tracked by GFP. The number of GFP positive foci per well normalized to cell area was calculated using Incucyte Analysis Software.

### Mucin-selective protease treatment of Calu-3 cells

Calu-3 cells overexpressing CRISPR activator components and MUC4- and NTG-targeting sgRNAs were seeded into 96-well plates, targeting 30% density for the next day. After allowing for attachment, media was changed to RPMI+2% FBS. 22 hours later, StcE mucinase diluted in RPMI+2% FBS was spiked into mucinase-treated wells to achieve a final concentration of 20nM mucinase. Vehicle conditions received RPMI+2% FBS to maintain equal well volumes. 2 hours later, the pseudovirus VSVdG-eGFP-CoV-2-S and control VSVdG-eGFP-RABV-G were added to achieve MOI 0.1, then centrifuged at 900xg at 30°C for 60 minutes. After spinfection, individual wells were imaged over time using an Incucyte System (Sartorius) in a 37°C incubator at 5% CO_2_. Images were taken of each well at 4x magnification, and infection was tracked by GFP. The number of GFP-positive foci per well was calculated using Incucyte Analysis Software.

### Single-cell RNA meta-analysis of TMPSSR2+ACE2+ ciliated lung epithelial cells

AUC values for transcripts for ciliated human lung epithelial cells co-expressing ACE2 and TMPSSR2 were obtained from published supplementary data from (Muus et al., 2021). AUC values for the top 100 hits based on MAGeCK rank for each screen that were present in the dataset were retrieved and sorted high to low for each screen in the screen comparison.

### Bulk RNA-seq analysis of mucin gene expression in response to SARS-CoV-2 infection

We identified previously reported RNA-seq datasets of diverse cells and tissues after SARS-CoV-2 infection in order to examine differential mucin gene expression. We derived Calu3-cell differential gene expression data and post-mortem human lung data from GSE147507 (Blanco-Melo et al., 2020), human alveolar organoid data and human lung graft data from GSE152586 (Katsura et al., 2020), mouse lung data from GSE154104 (Winkler et al., 2020), and hamster trachea and hamster lung data from GSE161200 (Hoagland et al., 2021). log2fold-change of transcript expression levels upon SARS-CoV-2 infection and false discovery rates (FDR) for the mucin genes were identified from each corresponding manuscript’s reported values. Missing values indicate no expression data was reported for that gene in the given dataset.

### RNA-seq analysis for COVID-19 clinical samples

Filtered feature-barcode matrices for 13 BALF samples (4 healthy controls, 9 individuals with COVID-19 were obtained from GSE145926 and GSM3660650 (NCBI). The standard preprocessing workflow in Seurat was followed. Samples were analyzed and further filtered down using quality control metrics to remove batch effects. After removing unwanted cells, each sample was normalized using ‘LogNormalize’ and integrated into one assay using Seurat v4. Linear transformation was applied to the data using ‘ScaleData’ prior to dimensional reduction. The top 2,000 variable genes were identified using the ‘FindVariableFeatures’ method and were used to perform PCA. UMAP was then performed using the top 50 principal components. Seurat uses a graph-based approach to cluster cells which was applied here using the ‘FindNeighbors’ and ‘FindClusters’ functions. Once each cell was assigned to a cluster, genetic markers KRT18, KRT19 and TPPP3 were used to identify epithelial cell clusters. The integrated assay was then subsetted to only include epithelial cells. The cells were regrouped based on the sample donor’s disease state. Gene expression data was extracted from the epithelial cell subset using ‘GetAssayData’ on its RNA assay. Finally, differentially expressed genes were identified between the healthy control cells and COVID19-infected cells using the ‘FindMarkers’ method. We calculated differential gene expression using the scDD R package (Korthauer, Keegan et al. 2019) with the hyperparameters a0 = 0.01, b0 = 0.01, mu0 = 0, s0 = 0.01, and alpha = 0.01.

## References

Almuttaqi, H., and Udalova, I.A. (2019). Advances and challenges in targeting IRF5, a key regulator of inflammation. FEBS J. 286, 1624–1637.

Arons, M.M., Hatfield, K.M., Reddy, S.C., Kimball, A., James, A., Jacobs, J.R., Taylor, J., Spicer, K., Bardossy, A.C., Oakley, L.P., et al. (2020). Presymptomatic SARS-CoV-2 Infections and Transmission in a Skilled Nursing Facility. N. Engl. J. Med. 382, 2081–2090.

Bagga, S., and Bouchard, M.J. (2014). Cell Cycle Regulation During Viral Infection. In Cell Cycle Control: Mechanisms and Protocols, E. Noguchi, and M.C. Gadaleta, eds. (New York, NY: Springer New York), pp. 165–227.

Baggen, J., Persoons, L., Vanstreels, E., Jansen, S., Van Looveren, D., Boeckx, B., Geudens, V., De Man, J., Jochmans, D., Wauters, J., et al. (2021). Genome-wide CRISPR screening identifies TMEM106B as a proviral host factor for SARS-CoV-2. Nat. Genet. 1–10.

Bailey, A.L., and Diamond, M.S. (2021). A Crisp(r) New Perspective on SARS-CoV-2 Biology. Cell 184, 15–17.

Balabanian, K., Levoye, A., Klemm, L., Lagane, B., Hermine, O., Harriague, J., Baleux, F., Arenzana-Seisdedos, F., and Bachelerie, F. (2008). Leukocyte analysis from WHIM syndrome patients reveals a pivotal role for GRK3 in CXCR4 signaling. J. Clin. Invest. 118, 1074–1084.

Bekisz, J., Baron, S., Balinsky, C., Morrow, A., and Zoon, K.C. (2010). Antiproliferative Properties of Type I and Type II Interferon. Pharmaceuticals 3, 994–1015.

Bennett, K.L., Modrell, B., Greenfield, B., Bartolazzi, A., Stamenkovic, I., Peach, R., Jackson, D.G., Spring, F., and Aruffo, A. (1995). Regulation of CD44 binding to hyaluronan by glycosylation of variably spliced exons. J. Cell Biol. 131, 1623–1633.

Bergelson, J.M. (2009). Intercellular junctional proteins as receptors and barriers to virus infection and spread. Cell Host Microbe 5, 517–521.

Biering, S.B., Van Dis, E., Wehri, E., Yamashiro, L.H., Nguyenla, X., Dugast-Darzacq, C., Graham, T.G.W., Stroumza, J.R., Golovkine, G.R., Roberts, A.W., et al. (2020). Screening a library of FDA-approved and bioactive compounds for antiviral activity against SARS-CoV-2.

Blanco-Melo, D., Nilsson-Payant, B.E., Liu, W.-C., Uhl, S., Hoagland, D., Møller, R., Jordan, T.X., Oishi, K., Panis, M., Sachs, D., et al. (2020). Imbalanced Host Response to SARS-CoV-2 Drives Development of COVID-19. Cell 181, 1036–1045.e9.

Bonifacino, J.S. (2014). Adaptor proteins involved in polarized sorting. J. Cell Biol. 204, 7–17.

Bouhaddou, M., Memon, D., Meyer, B., White, K.M., Rezelj, V.V., Correa Marrero, M., Polacco, B.J., Melnyk, J.E., Ulferts, S., Kaake, R.M., et al. (2020). The Global Phosphorylation Landscape of SARS-CoV-2 Infection. Cell 182, 685–712.e19.

Boyden, L.M., Atzmony, L., Hamilton, C., Zhou, J., Lim, Y.H., Hu, R., Pappas, J., Rabin, R., Ekstien, J., Hirsch, Y., et al. (2019). Recessive Mutations in AP1B1 Cause Ichthyosis, Deafness, and Photophobia. Am. J. Hum. Genet. 105, 1023–1029.

Brann, D.H., Tsukahara, T., Weinreb, C., Lipovsek, M., Van den Berge, K., Gong, B., Chance, R., Macaulay, I.C., Chou, H.-J., Fletcher, R.B., et al. (2020). Non-neuronal expression of SARS-CoV-2 entry genes in the olfactory system suggests mechanisms underlying COVID-19-associated anosmia. Sci Adv 6.

Button, B., Cai, L.-H., Ehre, C., Kesimer, M., Hill, D.B., Sheehan, J.K., Boucher, R.C., and Rubinstein, M. (2012). A Periciliary Brush Promotes the Lung Health by Separating the Mucus Layer from Airway Epithelia. Science 337, 937–941.

Carson, D.D. (2008). The Cytoplasmic Tail of MUC1: A Very Busy Place. Sci. Signal. 1, e35–pe35.

Castillon, G.A., Burriat-Couleru, P., Abegg, D., Criado Santos, N., and Watanabe, R. (2018). Clathrin and AP1 are required for apical sorting of glycosyl phosphatidyl inositol-anchored proteins in biosynthetic and recycling routes in Madin-Darby canine kidney cells. Traffic 19, 215–228.

Cawood, R., Harrison, S.M., Dove, B.K., Reed, M.L., and Hiscox, J.A. (2007). Cell cycle dependent nucleolar localization of the coronavirus nucleocapsid protein. Cell Cycle 6, 863–867.

Chen, G., Korfhagen, T.R., Xu, Y., Kitzmiller, J., Wert, S.E., Maeda, Y., Gregorieff, A., Clevers, H., and Whitsett, J.A. (2009). SPDEF is required for mouse pulmonary goblet cell differentiation and regulates a network of genes associated with mucus production. J. Clin. Invest. 119, 2914–2924.

Chen, Z., Wang, C., Feng, X., Nie, L., Tang, M., Zhang, H., Xiong, Y., Swisher, S.K., Srivastava, M., and Chen, J. (2021). Comprehensive analysis of the host-virus interactome of SARS-CoV-2. Biorxiv.

Clausen, T.M., Sandoval, D.R., Spliid, C.B., Pihl, J., Perrett, H.R., Painter, C.D., Narayanan, A., Majowicz, S.A., Kwong, E.M., McVicar, R.N., et al. (2020). SARS-CoV-2 Infection Depends on Cellular Heparan Sulfate and ACE2. Cell 183, 1043–1057.e15.

Cohen, M., Zhang, X.-Q., Senaati, H.P., Chen, H.-W., Varki, N.M., Schooley, R.T., and Gagneux, P. (2013). Influenza A penetrates host mucus by cleaving sialic acids with neuraminidase. Virol. J. 10, 321.

Cornillez-Ty, C.T., Liao, L., Yates, J.R., 3rd, Kuhn, P., and Buchmeier, M.J. (2009). Severe acute respiratory syndrome coronavirus nonstructural protein 2 interacts with a host protein complex involved in mitochondrial biogenesis and intracellular signaling. J. Virol. 83, 10314–10318.

da Costa, V.G., Moreli, M.L., and Saivish, M.V. (2020). The emergence of SARS, MERS and novel SARS-2 coronaviruses in the 21st century. Arch. Virol. 165, 1517–1526.

Cui, J., Li, F., and Shi, Z.-L. (2018). Origin and evolution of pathogenic coronaviruses. Nat. Rev. Microbiol. 17, 181–192.

Daniloski, Z., Jordan, T.X., Wessels, H.-H., Hoagland, D.A., Kasela, S., Legut, M., Maniatis, S., Mimitou, E.P., Lu, L., Geller, E., et al. (2021). Identification of Required Host Factors for SARS-CoV-2 Infection in Human Cells. Cell 184, 92–105.e16.

Datta, A., Sandilands, E., Mostov, K.E., and Bryant, D.M. (2017). Fibroblast-derived HGF drives acinar lung cancer cell polarization through integrin-dependent RhoA-ROCK1 inhibition. Cell. Signal. 40, 91–98.

Davy, C., and Doorbar, J. (2007). G2/M cell cycle arrest in the life cycle of viruses. Virology 368, 219–226.

DeDiego, M.L., Nieto-Torres, J.L., Regla-Nava, J.A., Jimenez-Guardeño, J.M., Fernandez-Delgado, R., Fett, C., Castaño-Rodriguez, C., Perlman, S., and Enjuanes, L. (2014). Inhibition of NF-κB-Mediated Inflammation in Severe Acute Respiratory Syndrome Coronavirus-Infected Mice Increases Survival. J. Virol. 88, 913–924.

Delaveris, C.S., Webster, E.R., Banik, S.M., Boxer, S.G., and Bertozzi, C.R. (2020). Membrane-tethered mucin-like polypeptides sterically inhibit binding and slow fusion kinetics of influenza A virus. Proc. Natl. Acad. Sci. U. S. A. 117, 12643–12650.

Desmyter, J., Melnick, J.L., and Rawls, W.E. (1968). Defectiveness of interferon production and of rubella virus interference in a line of African green monkey kidney cells (Vero). J. Virol. 2, 955–961.

Dieterle, M.E., Haslwanter, D., Bortz, R.H., 3rd, Wirchnianski, A.S., Lasso, G., Vergnolle, O., Abbasi, S.A., Fels, J.M., Laudermilch, E., Florez, C., et al. (2020). A Replication-Competent Vesicular Stomatitis Virus for Studies of SARS-CoV-2 Spike-Mediated Cell Entry and Its Inhibition. Cell Host Microbe 28, 486–496.e6.

Dyer, M.D., Murali, T.M., and Sobral, B.W. (2008). The Landscape of Human Proteins Interacting with Viruses and Other Pathogens. PLoS Pathog. 4, e32.

Ehre, C., Worthington, E.N., Liesman, R.M., Grubb, B.R., Barbier, D., O’Neal, W.K., Sallenave, J.-M., Pickles, R.J., and Boucher, R.C. (2012). Overexpressing mouse model demonstrates the protective role of Muc5ac in the lungs. Proc. Natl. Acad. Sci. U. S. A. 109, 16528–16533.

Ehrhardt, C., Rückle, A., Hrincius, E.R., Haasbach, E., Anhlan, D., Ahmann, K., Banning, C., Reiling, S.J., Kühn, J., Strobl, S., et al. (2013). The NF-κB inhibitor SC75741 efficiently blocks influenza virus propagation and confers a high barrier for development of viral resistance. Cell. Microbiol. 15, 1198–1211.

Fan, Y., Sanyal, S., and Bruzzone, R. (2018). Breaking bad: How viruses subvert the cell cycle. Front. Cell. Infect. Microbiol. 8, 396.

Flynn, R.A., Belk, J.A., Qi, Y., Yasumoto, Y., Schmitz, C.O., Mumbach, M.R., Limaye, A., Wei, J., Alfajaro, M.M., Parker, K.R., et al. (2020). Systematic discovery and functional interrogation of SARS-CoV-2 viral RNA-host protein interactions during infection. BioRxiv.

Fodoulian, L., Tuberosa, J., Rossier, D., Boillat, M., Kan, C., Pauli, V., Egervari, K., Lobrinus, J.A., Landis, B.N., Carleton, A., et al. (2020). SARS-CoV-2 Receptors and Entry Genes Are Expressed in the Human Olfactory Neuroepithelium and Brain. IScience 23, 101839.

Gasiorek, J.J., and Blank, V. (2015). Regulation and function of the NFE2 transcription factor in hematopoietic and non-hematopoietic cells. Cell. Mol. Life Sci. 72, 2323–2335.

Gordon, D.E., Jang, G.M., Bouhaddou, M., Xu, J., Obernier, K., White, K.M., O’Meara, M.J., Rezelj, V.V., Guo, J.Z., Swaney, D.L., et al. (2020). A SARS-CoV-2 protein interaction map reveals targets for drug repurposing. Nature 583, 459–468.

Guan, W.-J., Chen, R.-C., and Zhong, N.-S. (2020). Strategies for the prevention and management of coronavirus disease 2019. Eur. Respir. J. 55.

Hartenian, E., Nandakumar, D., Lari, A., Ly, M., Tucker, J.M., and Glaunsinger, B.A. (2020). The molecular virology of coronaviruses. J. Biol. Chem. 295, 12910–12934.

Hase, K., Nakatsu, F., Ohmae, M., Sugihara, K., Shioda, N., Takahashi, D., Obata, Y., Furusawa, Y., Fujimura, Y., Yamashita, T., et al. (2013). AP-1B-mediated protein sorting regulates polarity and proliferation of intestinal epithelial cells in mice. Gastroenterology 145, 625–635.

Hasegawa, M., Takahashi, H., Rajabi, H., Alam, M., Suzuki, Y., Yin, L., Tagde, A., Maeda, T., Hiraki, M., Sukhatme, V.P., et al. (2016). Functional interactions of the cystine/glutamate antiporter, CD44v and MUC1-C oncoprotein in triple-negative breast cancer cells. Oncotarget 7, 11756–11769.

Haston, C.K., Cory, S., Lafontaine, L., Dorion, G., and Hallett, M.T. (2006). Strain-dependent pulmonary gene expression profiles of a cystic fibrosis mouse model. Physiol. Genomics 25, 336–345.

Hattrup, C.L., and Gendler, S.J. (2008). Structure and function of the cell surface (tethered) mucins. Annu. Rev. Physiol. 70, 431–457.

He, J., Cai, S., Feng, H., Cai, B., Lin, L., Mai, Y., Fan, Y., Zhu, A., Huang, H., Shi, J., et al. (2020). Single-cell analysis reveals bronchoalveolar epithelial dysfunction in COVID-19 patients. Protein Cell 11, 680–687.

Hoagland, D.A., Møller, R., Uhl, S.A., Oishi, K., Frere, J., Golynker, I., Horiuchi, S., Panis, M., Blanco-Melo, D., Sachs, D., et al. (2021). Leveraging the antiviral type I interferon system as a first line of defense against SARS-CoV-2 pathogenicity. Immunity 54, 557–570.e5.

Hoffmann, M., Kleine-Weber, H., Schroeder, S., Krüger, N., Herrler, T., Erichsen, S., Schiergens, T.S., Herrler, G., Wu, N.-H., Nitsche, A., et al. (2020). SARS-CoV-2 Cell Entry Depends on ACE2 and TMPRSS2 and Is Blocked by a Clinically Proven Protease Inhibitor. Cell 181, 271–280.e8.

Honda, R., Lowe, E.D., Dubinina, E., Skamnaki, V., Cook, A., Brown, N.R., and Johnson, L.N. (2005). The structure of cyclin E1/CDK2: implications for CDK2 activation and CDK2-independent roles. EMBO J. 24, 452–463.

Honke, N., Shaabani, N., Zhang, D.-E., Hardt, C., and Lang, K.S. (2016). Multiple functions of USP18. Cell Death Dis. 7, e2444.

Hsu, P.D., Lander, E.S., and Zhang, F. (2014). Development and applications of CRISPR-Cas9 for genome engineering. Cell 157, 1262–1278.

Iverson, E., Griswold, K., Song, D., Gagliardi, T.B., Hamidzadeh, K., Kesimer, M., Sinha, S., Perry, M., Duncan, G.A., and Scull, M.A. (2021). Membrane-Tethered Mucin 1 is Stimulated by Interferon in Multiple Cell Types and Antagonizes Influenza A Virus Infection in Human Airway Epithelium.

Jefferies, C.A. (2019). Regulating IRFs in IFN Driven Disease. Front. Immunol. 10, 325.

Jia, X., Weber, E., Tokarev, A., Lewinski, M., Rizk, M., Suarez, M., Guatelli, J., and Xiong, Y. (2014). Structural basis of HIV-1 Vpu-mediated BST2 antagonism via hijacking of the clathrin adaptor protein complex 1. Elife 3, e02362.

Joung, J., Konermann, S., Gootenberg, J.S., Abudayyeh, O.O., Platt, R.J., Brigham, M.D., Sanjana, N.E., and Zhang, F. (2017). Genome-scale CRISPR-Cas9 knockout and transcriptional activation screening. Nat. Protoc. 12, 828–863.

Katsura, H., Sontake, V., Tata, A., Kobayashi, Y., Edwards, C.E., Heaton, B.E., Konkimalla, A., Asakura, T., Mikami, Y., Fritch, E.J., et al. (2020). Human Lung Stem Cell-Based Alveolospheres Provide Insights into SARS-CoV-2-Mediated Interferon Responses and Pneumocyte Dysfunction. Cell Stem Cell 27, 890–904.e8.

Katze, M.G., He, Y., and Gale, M., Jr (2002). Viruses and interferon: a fight for supremacy. Nat. Rev. Immunol. 2, 675–687.

Kesimer, M., Ehre, C., Burns, K.A., Davis, C.W., Sheehan, J.K., and Pickles, R.J. (2013). Molecular organization of the mucins and glycocalyx underlying mucus transport over mucosal surfaces of the airways. Mucosal Immunol. 6, 379–392.

Kircheis, R., Haasbach, E., Lueftenegger, D., Heyken, W.T., Ocker, M., and Planz, O. (2020). NF-κB Pathway as a Potential Target for Treatment of Critical Stage COVID-19 Patients. Front. Immunol. 11, 598444.

Koch, J., Uckeley, Z.M., Doldan, P., Stanifer, M., Boulant, S., and Lozach, P.-Y. (2020). Host Cell Proteases Drive Early or Late SARS-CoV-2 Penetration.

Konermann, S., Brigham, M.D., Trevino, A.E., Joung, J., Abudayyeh, O.O., Barcena, C., Hsu, P.D., Habib, N., Gootenberg, J.S., Nishimasu, H., et al. (2015). Genome-scale transcriptional activation by an engineered CRISPR-Cas9 complex. Nature 517, 583–588.

Kost-Alimova, M., Sidhom, E.-H., Satyam, A., Chamberlain, B.T., Dvela-Levitt, M., Melanson, M., Alper, S.L., Santos, J., Gutierrez, J., Subramanian, A., et al. (2020). A High-Content Screen for Mucin-1-Reducing Compounds Identifies Fostamatinib as a Candidate for Rapid Repurposing for Acute Lung Injury. Cell Reports Medicine 1, 100137.

Krishnan, M.N., Ng, A., Sukumaran, B., Gilfoy, F.D., Uchil, P.D., Sultana, H., Brass, A.L., Adametz, R., Tsui, M., Qian, F., et al. (2008). RNA interference screen for human genes associated with West Nile virus infection. Nature 455, 242–245.

Kumar, N., Xin, Z.-T., Liang, Y., Ly, H., and Liang, Y. (2008). NF-kappaB signaling differentially regulates influenza virus RNA synthesis. J. Virol. 82, 9880–9889.

Lee, I.T., Nakayama, T., Wu, C.-T., Goltsev, Y., Jiang, S., Gall, P.A., Liao, C.-K., Shih, L.-C., Schürch, C.M., McIlwain, D.R., et al. (2020). ACE2 localizes to the respiratory cilia and is not increased by ACE inhibitors or ARBs. Nat. Commun. 11, 5453.

Lee, R.J., Xiong, G., Kofonow, J.M., Chen, B., Lysenko, A., Jiang, P., Abraham, V., Doghramji, L., Adappa, N.D., Palmer, J.N., et al. (2012). T2R38 taste receptor polymorphisms underlie susceptibility to upper respiratory infection. J. Clin. Invest. 122, 4145–4159.

Li, X., Commane, M., Nie, H., Hua, X., Chatterjee-Kishore, M., Wald, D., Haag, M., and Stark, G.R. (2000). Act1, an NF-kappa B-activating protein. Proc. Natl. Acad. Sci. U. S. A. 97, 10489–10493.

Liao, M., Liu, Y., Yuan, J., Wen, Y., Xu, G., Zhao, J., Chen, L., Li, J., Wang, X., Wang, F., et al. (2020). The landscape of lung bronchoalveolar immune cells in COVID-19 revealed by singlecell RNA sequencing (medRxiv).

Lillehoj, E.P., Kato, K., Lu, W., and Kim, K.C. (2013). Cellular and molecular biology of airway mucins. Int. Rev. Cell Mol. Biol. 303, 139–202.

Liu, Y., Lv, J., Liu, J., Li, M., Xie, J., Lv, Q., Deng, W., Zhou, N., Zhou, Y., Song, J., et al. (2020). Mucus production stimulated by IFN-AhR signaling triggers hypoxia of COVID-19. Cell Res. 30, 1078–1087.

Lu, W., Hisatsune, A., Koga, T., Kato, K., Kuwahara, I., Lillehoj, E.P., Chen, W., Cross, A.S., Gendler, S.J., Gewirtz, A.T., et al. (2006). Cutting edge: enhanced pulmonary clearance of Pseudomonas aeruginosa by Muc1 knockout mice. J. Immunol. 176, 3890–3894.

Lu, W., Liu, X., Wang, T., Liu, F., Zhu, A., Lin, Y., Luo, J., Ye, F., He, J., Zhao, J., et al. (2021). Elevated MUC1 and MUC5AC mucin protein levels in airway mucus of critical ill COVID-19 patients. J. Med. Virol. 93, 582–584.

Lukassen, S., Chua, R.L., Trefzer, T., Kahn, N.C., Schneider, M.A., Muley, T., Winter, H., Meister, M., Veith, C., Boots, A.W., et al. (2020). SARS-CoV-2 receptor ACE2 and TMPRSS2 are primarily expressed in bronchial transient secretory cells. EMBO J. 39, e105114.

Ma, J., Rubin, B.K., and Voynow, J.A. (2018). Mucins, Mucus, and Goblet Cells. Chest 154, 169–176.

Ma, S., Meng, Z., Chen, R., and Guan, K.-L. (2019). The Hippo Pathway: Biology and Pathophysiology. Annu. Rev. Biochem. 88, 577–604.

Malaker, S.A., Pedram, K., Ferracane, M.J., Bensing, B.A., Krishnan, V., Pett, C., Yu, J., Woods, E.C., Kramer, J.R., Westerlind, U., et al. (2019). The mucin-selective protease StcE enables molecular and functional analysis of human cancer-associated mucins. Proc. Natl. Acad. Sci. U. S. A. 116, 7278–7287.

McAuley, J.L., Corcilius, L., Tan, H.-X., Payne, R.J., McGuckin, M.A., and Brown, L.E. (2017). The cell surface mucin MUC1 limits the severity of influenza A virus infection. Mucosal Immunol. 10, 1581–1593.

Mizutani, T., Fukushi, S., Iizuka, D., Inanami, O., Kuwabara, M., Takashima, H., Yanagawa, H., Saijo, M., Kurane, I., and Morikawa, S. (2006). Inhibition of cell proliferation by SARS-CoV infection in Vero E6 cells. FEMS Immunol. Med. Microbiol. 46, 236–243.

Mosca, J.D., and Pitha, P.M. (1986). Transcriptional and posttranscriptional regulation of exogenous human beta interferon gene in simian cells defective in interferon synthesis. Mol. Cell. Biol. 6, 2279–2283.

Mothes, W., Sherer, N.M., Jin, J., and Zhong, P. (2010). Virus Cell-to-Cell Transmission. J. Virol. 84, 8360–8368.

Muus, C., Luecken, M.D., Eraslan, G., Sikkema, L., Waghray, A., Heimberg, G., Kobayashi, Y., Vaishnav, E.D., Subramanian, A., Smillie, C., et al. (2021). Single-cell meta-analysis of SARS-CoV-2 entry genes across tissues and demographics. Nat. Med. 27, 546–559.

Nakashima, T., Yokoyama, A., Ohnishi, H., Hamada, H., Ishikawa, N., Haruta, Y., Hattori, N., Tanigawa, K., and Kohno, N. (2008). Circulating KL-6/MUC1 as an independent predictor for disseminated intravascular coagulation in acute respiratory distress syndrome. J. Intern. Med. 263, 432–439.

Navarro, Negredo P., Edgar, J.R., Wrobel, A.G., Zaccai, N.R., Antrobus, R., Owen, D.J., and Robinson, M.S. (2017). Contribution of the clathrin adaptor AP-1 subunit μ1 to acidic cluster protein sorting. J. Cell Biol. 216, 2927–2943.

Nguyen, L., McCord, K.A., Bui, D.T., Bouwman, K.M., Kitova, E.N., Kumawat, D., Daskhan, G.C., Tomris, I., Han, L., Chopra, P., et al. (2021). Sialic acid-Dependent Binding and Viral Entry of SARS-CoV-2.

Orchard, R.C., Wilen, C.B., Doench, J.G., Baldridge, M.T., McCune, B.T., Lee, Y.-C.J., Lee, S., Pruett-Miller, S.M., Nelson, C.A., Fremont, D.H., et al. (2016). Discovery of a proteinaceous cellular receptor for a norovirus. Science 353, 933–936.

Ou, T., Mou, H., Zhang, L., Ojha, A., Choe, H., and Farzan, M. (2021). Hydroxychloroquine-mediated inhibition of SARS-CoV-2 entry is attenuated by TMPRSS2. PLoS Pathog. 17, e1009212.

Park, M.H., and Hong, J.T. (2016). Roles of NF-κB in Cancer and Inflammatory Diseases and Their Therapeutic Approaches. Cells 5.

Pfaender, S., Mar, K.B., Michailidis, E., Kratzel, A., Boys, I.N., V’kovski, P., Fan, W., Kelly, J.N., Hirt, D., Ebert, N., et al. (2020). LY6E impairs coronavirus fusion and confers immune control of viral disease. Nature Microbiology 5, 1330–1339.

Pinto, R., Herold, S., Cakarova, L., Hoegner, K., Lohmeyer, J., Planz, O., and Pleschka, S. (2011). Inhibition of influenza virus-induced NF-kappaB and Raf/MEK/ERK activation can reduce both virus titers and cytokine expression simultaneously in vitro and in vivo. Antiviral Res. 92, 45–56.

Plante, J.A., Plante, K.S., Gralinski, L.E., Beall, A., Ferris, M.T., Bottomly, D., Green, R., McWeeney, S.K., Heise, M.T., Baric, R.S., et al. (2020). Mucin 4 Protects Female Mice from Coronavirus Pathogenesis.

Plouffe, S.W., Meng, Z., Lin, K.C., Lin, B., Hong, A.W., Chun, J.V., and Guan, K.-L. (2016). Characterization of Hippo Pathway Components by Gene Inactivation. Mol. Cell 64, 993–1008.

Poppe, M., Wittig, S., Jurida, L., Bartkuhn, M., Wilhelm, J., Müller, H., Beuerlein, K., Karl, N., Bhuju, S., Ziebuhr, J., et al. (2017). The NF-κB-dependent and -independent transcriptome and chromatin landscapes of human coronavirus 229E-infected cells. PLoS Pathog. 13, e1006286.

Ravindra, N.G., Alfajaro, M.M., Gasque, V., Huston, N.C., Wan, H., Szigeti-Buck, K., Yasumoto, Y., Greaney, A.M., Habet, V., Chow, R.D., et al. (2021). Single-cell longitudinal analysis of SARS-CoV-2 infection in human airway epithelium identifies target cells, alterations in gene expression, and cell state changes. PLoS Biol. 19, e3001143.

Reay, W.R., Geaghan, M.P., Cairns, M.J., and 23andMe Research Team (2021). Genome-wide meta-analysis of pneumonia suggests a role for mucin biology and provides novel drug repurposing opportunities (medRxiv).

Sahajpal, N.S., Lai, C.-Y.J., Hastie, A., Mondal, A.K., Dehkordi, S.R., van der Made, C., Fedrigo, O., Al-Ajli, F., Jalnapurkar, S., Kanagal-Shamanna, R., et al. (2021). Host genome analysis of structural variations by Optical Genome Mapping provides clinically valuable insights into genes implicated in critical immune, viral infection, and viral replication pathways in patients with severe COVID-19. MedRxiv 2021.01.05.21249190.

Sajuthi, S.P., DeFord, P., Li, Y., Jackson, N.D., Montgomery, M.T., Everman, J.L., Rios, C.L., Pruesse, E., Nolin, J.D., Plender, E.G., et al. (2020). Type 2 and interferon inflammation regulate SARS-CoV-2 entry factor expression in the airway epithelium. Nat. Commun. 11, 1–18.

Salas, A., Pardo-Seco, J., Cebey-López, M., Gómez-Carballa, A., Obando-Pacheco, P., Rivero-Calle, I., Currás-Tuala, M.-J., Amigo, J., Gómez-Rial, J., Martinón-Torres, F., et al. (2017). Whole Exome Sequencing reveals new candidate genes in host genomic susceptibility to Respiratory Syncytial Virus Disease. Sci. Rep. 7, 15888.

Sanson, K.R., Hanna, R.E., Hegde, M., Donovan, K.F., Strand, C., Sullender, M.E., Vaimberg, E.W., Goodale, A., Root, D.E., Piccioni, F., et al. (2018). Optimized libraries for CRISPR-Cas9 genetic screens with multiple modalities. Nat. Commun. 9, 5416.

Santoro, M.G., Rossi, A., and Amici, C. (2003). NF-kappaB and virus infection: who controls whom. EMBO J. 22, 2552–2560.

Schmitz, M.L., Kracht, M., and Saul, V.V. (2014). The intricate interplay between RNA viruses and NF-κB. Biochim. Biophys. Acta 1843, 2754–2764.

Schmolke, M., Viemann, D., Roth, J., and Ludwig, S. (2009). Essential Impact of NF-κB Signaling on the H5N1 Influenza A Virus-Induced Transcriptome. The Journal of Immunology 183, 5180–5189.

Schneider, W.M., Luna, J.M., Hoffmann, H.-H., Sánchez-Rivera, F.J., Leal, A.A., Ashbrook, A.W., Le Pen, J., Ricardo-Lax, I., Michailidis, E., Peace, A., et al. (2021). Genome-Scale Identification of SARS-CoV-2 and Pan-coronavirus Host Factor Networks. Cell 184, 120–132.e14.

Shah, A.S., Ben-Shahar, Y., Moninger, T.O., Kline, J.N., and Welsh, M.J. (2009). Motile cilia of human airway epithelia are chemosensory. Science 325, 1131–1134.

Strich, J.R. (2020). Fostamatinib for Hospitalized Adults With COVID-19.

Su, M., Chen, Y., Qi, S., Shi, D., Feng, L., and Sun, D. (2020). A mini-review on cell cycle regulation of Coronavirus infection. Front. Vet. Sci. 7, 586826.

Sumpter, R., Jr, Loo, Y.-M., Foy, E., Li, K., Yoneyama, M., Fujita, T., Lemon, S.M., and Gale, M., Jr (2005). Regulating intracellular antiviral defense and permissiveness to hepatitis C virus RNA replication through a cellular RNA helicase, RIG-I. J. Virol. 79, 2689–2699.

Sungnak, W., Huang, N., Bécavin, C., Berg, M., Queen, R., Litvinukova, M., Talavera-López, C., Maatz, H., Reichart, D., Sampaziotis, F., et al. (2020). SARS-CoV-2 entry factors are highly expressed in nasal epithelial cells together with innate immune genes. Nat. Med. 26, 681–687.

Surjit, M., Liu, B., Chow, V.T.K., and Lal, S.K. (2006). The nucleocapsid protein of severe acute respiratory syndrome-coronavirus inhibits the activity of cyclin-cyclin-dependent kinase complex and blocks S phase progression in mammalian cells. J. Biol. Chem. 281, 10669–10681.

Tabassum, N., Zhang, H., and Stebbing, J. (2020). Repurposing Fostamatinib to Combat SARS-CoV-2-Induced Acute Lung Injury. Cell Rep Med 1, 100145.

Takahashi, D., Hase, K., Kimura, S., Nakatsu, F., Ohmae, M., Mandai, Y., Sato, T., Date, Y., Ebisawa, M., Kato, T., et al. (2011). The epithelia-specific membrane trafficking factor AP-1B controls gut immune homeostasis in mice. Gastroenterology 141, 621–632.

Taniguchi, K., and Karin, M. (2018). NF-κB, inflammation, immunity and cancer: coming of age. Nat. Rev. Immunol. 18, 309–324.

Taylor, M.P., Koyuncu, O.O., and Enquist, L.W. (2011). Subversion of the actin cytoskeleton during viral infection. Nat. Rev. Microbiol. 9, 427–439.

Thair, S.A., He, Y.D., Hasin-Brumshtein, Y., Sakaram, S., Pandya, R., Toh, J., Rawling, D., Remmel, M., Coyle, S., Dalekos, G.N., et al. (2021). Transcriptomic similarities and differences in host response between SARS-CoV-2 and other viral infections. IScience 24, 101947.

Trougakos, I.P., Stamatelopoulos, K., Terpos, E., Tsitsilonis, O.E., Aivalioti, E., Paraskevis, D., Kastritis, E., Pavlakis, G.N., and Dimopoulos, M.A. (2021). Insights to SARS-CoV-2 life cycle, pathophysiology, and rationalized treatments that target COVID-19 clinical complications. J. Biomed. Sci. 28, 9.

Vestbo, J. (2002). Epidemiological studies in mucus hypersecretion. Novartis Found. Symp. 248, 3-12; discussion 12–9, 277-282.

V’kovski, P., Kratzel, A., Steiner, S., Stalder, H., and Thiel, V. (2020). Coronavirus biology and replication: implications for SARS-CoV-2. Nat. Rev. Microbiol. 19, 155–170.

Wagner, C.E., Wheeler, K.M., and Ribbeck, K. (2018). Mucins and Their Role in Shaping the Functions of Mucus Barriers. Annu. Rev. Cell Dev. Biol. 34, 189–215.

Wang, R., Simoneau, C.R., Kulsuptrakul, J., Bouhaddou, M., Travisano, K.A., Hayashi, J.M., Carlson-Stevermer, J., Zengel, J.R., Richards, C.M., Fozouni, P., et al. (2021). Genetic Screens Identify Host Factors for SARS-CoV-2 and Common Cold Coronaviruses. Cell 184, 106–119.e14.

Wang, Y., Yuan, S., Jia, X., Ge, Y., Ling, T., Nie, M., Lan, X., Chen, S., and Xu, A. (2019). Mitochondria-localised ZNFX1 functions as a dsRNA sensor to initiate antiviral responses through MAVS. Nat. Cell Biol. 21, 1346–1356.

Wei, J., Alfajaro, M.M., DeWeirdt, P.C., Hanna, R.E., Lu-Culligan, W.J., Cai, W.L., Strine, M.S., Zhang, S.-M., Graziano, V.R., Schmitz, C.O., et al. (2021). Genome-wide CRISPR Screens Reveal Host Factors Critical for SARS-CoV-2 Infection. Cell 184, 76–91.e13.

Wen, Z., Zhang, Y., Lin, Z., Shi, K., and Jiu, Y. (2020). Cytoskeleton—a crucial key in host cell for coronavirus infection. J. Mol. Cell Biol. 12, 968–979.

Winkler, E.S., Bailey, A.L., Kafai, N.M., Nair, S., McCune, B.T., Yu, J., Fox, J.M., Chen, R.E., Earnest, J.T., Keeler, S.P., et al. (2020). SARS-CoV-2 infection of human ACE2-transgenic mice causes severe lung inflammation and impaired function. Nat. Immunol. 21, 1327–1335.

Yang, H., Lyu, Y., and Hou, F. (2021). SARS-CoV-2 infection and the antiviral innate immune response. J. Mol. Cell Biol. 12, 963–967.

Yang, X., Yu, Y., Xu, J., Shu, H., Xia, J., Liu, H., Wu, Y., Zhang, L., Yu, Z., Fang, M., et al. (2020). Clinical course and outcomes of critically ill patients with SARS-CoV-2 pneumonia in Wuhan, China: a single-centered, retrospective, observational study. The Lancet Respiratory Medicine 8, 475–481.

Yuan, X., Wu, J., Shan, Y., Yao, Z., Dong, B., Chen, B., Zhao, Z., Wang, S., Chen, J., and Cong, Y. (2006). SARS coronavirus 7a protein blocks cell cycle progression at G0/G1 phase via the cyclin D3/pRb pathway. Virology 346, 74–85.

Yuan, X., Yao, Z., Wu, J., Zhou, Y., Shan, Y., Dong, B., Zhao, Z., Hua, P., Chen, J., and Cong, Y. (2007). G1 phase cell cycle arrest induced by SARS-CoV 3a protein via the cyclin D3/pRb pathway. Am. J. Respir. Cell Mol. Biol. 37, 9–19.

Zanin, M., Baviskar, P., Webster, R., and Webby, R. (2016). The Interaction between Respiratory Pathogens and Mucus. Cell Host Microbe 19, 159–168.

Zazhytska, M., Kodra, A., Hoagland, D.A., Fullard, J.F., Shayya, H., Omer, A., Firestein, S., Gong, Q., Canoll, P.D., Goldman, J.E., et al. (2021). Disruption of nuclear architecture as a cause of COVID-19 induced anosmia.

Zhang, Q., Chen, C.Z., Swaroop, M., Xu, M., Wang, L., Lee, J., Wang, A.Q., Pradhan, M., Hagen, N., Chen, L., et al. (2020). Heparan sulfate assists SARS-CoV-2 in cell entry and can be targeted by approved drugs in vitro. Cell Discovery 6, 1–14.

Zhu, Y., Feng, F., Hu, G., Wang, Y., Yu, Y., Zhu, Y., Xu, W., Cai, X., Sun, Z., Han, W., et al. (2021). A genome-wide CRISPR screen identifies host factors that regulate SARS-CoV-2 entry. Nat. Commun. 12, 1–11.

